# Drosophila Hedgehog can act as a morphogen in the absence of regulated Ci processing

**DOI:** 10.1101/2020.06.23.167585

**Authors:** Jamie C. Little, Elisa Garcia-Garcia, Amanda Sul, Daniel Kalderon

## Abstract

Extracellular Hedgehog (Hh) proteins induce transcriptional changes in target cells by inhibiting the proteolytic processing of full-length Drosophila Ci or mammalian Gli proteins to nuclear transcriptional repressors and by activating the full-length proteins, which are otherwise held inactive by cytoplasmic binding partners and subject to accelerated degradation following activation. We used Ci variants expressed at physiological levels to investigate the contributions of these mechanisms to dose-dependent Hh signaling at the anteroposterior (AP) border of Drosophila wing imaginal discs. Ci variants that cannot be processed supported a normal pattern of graded target gene activation and the development of adults with normal wing morphology when supplemented by constitutive Ci repressor, showing that Hh can signal normally in the absence of regulated processing. The full-length Ci-155 protein profile of these variants revealed a linear gradient of Hh-stimulated degradation, allowing derivation of a spatial profile of inhibition of processing of normal C-155 by Hh. The processing-resistant Ci variants were also significantly activated in the absence of Hh by elimination of Cos2, acting through association with the CORD domain of Ci, or PKA, revealing separate inhibitory roles of these two components in addition to their well-established roles in promoting Ci-155 processing.

## Introduction

Hedgehog (Hh) signaling proteins guide development and help maintain adult tissue homeostasis in both invertebrates and vertebrates (Hui and Angers, 2011; Ingham and McMahon, 2001; Petrova and Joyner, 2014). Aberrant Hh protein production, distribution, and responses are common causes of developmental birth defects and cancer including holoprosencephaly, limb and digit abnormalities, medulloblastoma and basal cell carcinoma (Anderson et al., 2012; Cortes et al., 2019; Ng and Curran, 2011; Pak and Segal, 2016; Petrova and Joyner, 2014; Sasai et al., 2019). Understanding the basic molecular mechanisms of Hh communication is the first step in combating these various Hh related disorders. Many conserved Hh components were initially identified in *Drosophila melanogaster* and then found to have a mammalian ortholog, including the key transducing protein Smoothened (Smo), which is now the target of several anti-cancer drugs (Cortes et al., 2019; Ng and Curran, 2011; Pak and Segal, 2016). There are also differences between Drosophila and mammalian Hh signal transduction but neither pathway is fully understood (Briscoe and Therond, 2013; Huangfu and Anderson, 2006; Kong et al., 2019; Lee et al., 2016; Liu, 2019). Hence, it is important to understand the precise molecular mechanisms involved in the pathway in Drosophila, which allows complex genetic tests under physiological conditions.

In flies, multiple Hh signaling components interact to elicit the induction and de-repression of Hh target genes through Cubitus Interruptus (Ci), the singular transcription factor of the pathway (Dominguez et al., 1996; Methot and Basler, 2001; Xiong et al., 2015). Notably, Hh can act as a morphogen that signals to Ci to transcribe different Hh target gene products depending on how much ligand is present at the cell membrane. In third instar larval Drosophila wing discs, Hh is expressed in posterior compartment cells and Ci is expressed only in anterior cells, so that Hh signals to a band of anterior cells at the anterior-posterior (AP) border with declining strength from posterior to anterior (Blair, 2003; Lawrence and Struhl, 1996). Within this territory, full-length Ci (Ci-155) is activated to induce *decapentaplegic* (*dpp*) in a broad region, *patched* (*ptc* or a *ptc-lacZ* transcriptional reporter) in a gradient, and *engrailed (en*) only in the cells closest to the source of Hh (Blair, 2003; Vervoort, 2000). In order for Ci to perform these functions, multiple upstream signaling proteins dynamically regulate Ci proteolytic processing, activation, and degradation.

In the absence of Hh, the receptor Patched (Ptc) actively inhibits the actions of another transmembrane protein Smoothened (Smo), which is kept at low levels and mainly associated with internal vesicles (Strigini and Cohen, 1997; Zhao et al., 2007). Costal2 (Cos2), complexed to Fused (Fu), acts as a scaffold for Protein Kinase A (PKA), Glycogen Synthase Kinase-3 (GSK3), and Casein Kinase-1 (CK1) to associate with C-155 and phosphorylate Ci-155 at a series of clustered sites (Ranieri et al., 2014; Zhang et al., 2005). This creates a binding site for Slimb, the substrate recognition component of the SCF ubiquitin ligase complex, which promotes the ubiquitination and subsequent partial proteolysis of full length Ci-155 by the proteasome to a shortened repressor form (Ci-75) lacking the C-terminal half of Ci-155, including its transcriptional activation domain (Aza-Blanc et al., 1997; Jia et al., 2005; Jiang, 2006; Smelkinson and Kalderon, 2006; Smelkinson et al., 2007).

Hh binding to Ptc leads to Smo activation in a process that includes phosphorylation at clustered C-terminal sites by PKA, CK1and GRK2, accumulation at the plasma membrane and a change in conformation or oligomerization (Kalderon, 2008; Maier et al., 2014; Zhao et al., 2007). Activation alters binding of Smo to Cos2-Fu complexes, with two important consequences. First, Ci-155 processing is inhibited, reportedly due to titration of Cos2 complexes away from Ci-155 and perhaps also to partial dissociation of those complexes (Li et al., 2014; Ranieri et al., 2014; Zhang et al., 2005). Second, Fu monomers are brought together to cross-phosphorylate and activate Fu protein kinase activity (Shi et al., 2011; Zhang et al., 2011; Zhou and Kalderon, 2011). Activated Fu protein kinase is necessary for the full activation of Ci-155, which is otherwise maintained in an inactive cytoplasmic form through direct associations with Suppressor of fused (Su(fu)) and Cos2 (Forbes et al., 1993; Ohlmeyer and Kalderon, 1998; Preat et al., 1993). Fu protein associations, but not kinase activity, are required for Ci-155 processing, so the role of Fu kinase activity in Ci-155 activation was studied in isolation by using point mutations in the kinase domain that only affect activation (Therond et al., 1996; Zadorozny et al., 2015). Active Ci-155 and Ci-75 repressor share the same zinc finger DNA-binding domain and have opposing transcriptional effects on the regulation of Hh target gene induction. Nevertheless, individual target genes have different sensitivities to Ci repressor and activator depending on the arrangement of Ci binding sites and the tonic influence of other transcription factors (Biehs et al., 2010; Methot and Basler, 2001; Muller and Basler, 2000; Parker et al., 2011). Thus, for example, repression is essential to silence *dpp* but not *ptc* or *en* in anterior cells away from the wing disc AP border. Activation of Ci-155 is critical to Hedgehog (Hh) signaling and is barely understood (Ohlmeyer and Kalderon, 1998; Zhou and Kalderon, 2011). It is also not clear to what extent regulation of Ci-155 activation and Ci-155 processing, as well as Hh-stimulated Ci-155 degradation, contribute independently to Hh morphogen action or whether these Hh-stimulated changes are inter-dependent. To address these issues we set out to study how processing-resistant Ci variants affected Ci-155 protein levels and activity.

The properties of Ci variants in wing discs have been investigated with some success using convenient conditions of non-physiological *GAL4*-responsive *UAS*-driven transgene expression levels, often in posterior wing disc cells to exclude endogenous Ci (Jia et al., 2005; Smelkinson et al., 2007). However, we previously found that Ci-155 activation, in contrast to Ci-155 processing, cannot be studied reliably in this way (Garcia-Garcia et al., 2017). Specifically, *ci*-null animals are very rarely rescued to adulthood using different combinations of *ci-Gal4* and *UAS-Ci* transgenes at a variety of temperatures and anterior En expression at the AP border was not rescued in *ci*-null clones by *UAS-Ci* expressed with the commonly used wing disc driver *C765-Gal4*, with transgene expression alone sometimes eliciting a dominant-negative effect on Hh target gene expression (Garcia-Garcia et al., 2017). We therefore developed genomic *ci* transgenes (Garcia-Garcia et al., 2017) and an efficient CRISPR strategy to directly alter the *ci* gene itself in order to study the full range of variant Ci activities under strictly physiological conditions.

In this study, we examined several processing-resistant Ci variants and found that Ci-155 protein levels were elevated to maximal levels in anterior wing disc cells away from the AP border, confirming full inhibition of Ci-155 processing. At the AP border there was a prominent graded decline of Ci-155 protein from anterior to posterior for those variants fully activated by Hh, providing the clearest image yet of graded Hh-stimulated Ci-155 degradation. Remarkably, the pattern and strength of *ptc-lacZ* and En induction in those wing discs was normal and processing-resistant Ci variants were also found to support the development of adults with normal wing patterning, provided a constitutive source of Ci repressor was provided. Thus, Ci can mediate normal Hh morphogen action in wing discs in the complete absence of regulated processing. We also used these processing-resistant Ci variants to study the potential roles of PKA and Cos2 in regulating Ci-155 activity in isolation from their well-established role in Ci-155 processing. We found that, in the absence of Hh, PKA inhibits Ci activity independent of the phosphorylation sites that regulate processing and that Cos2 inhibits Ci-155 activity by binding to the CORD region on Ci-155.

## Results

### Functional transgenes and CRISPR ci alleles

We developed strategies to study Ci expressed at physiological levels from “genomic *ci*” transgenes (*gCi*) and CRISPR-engineered *ci* alleles (*crCi*). The former strategy used a 16kb genomic region of *ci* that included upstream and downstream regulatory regions (Fig. 1A) previously used to rescue *ci* nulls using a second chromosome P-element insertion (Methot and Basler, 1999), inserted into an *att* site on the third chromosome (Garcia-Garcia et al., 2017). A *gCi-WT* transgene was readily able to rescue homozygous *ci* null (*ci^94^*) animals to adulthood, with normal morphology, and behaved almost like a normal *ci* allele but with marginally lower *ci* expression and activity in wing discs (Fig. 1E). We created *gCi* variants using this strategy.

**Figure 1.**
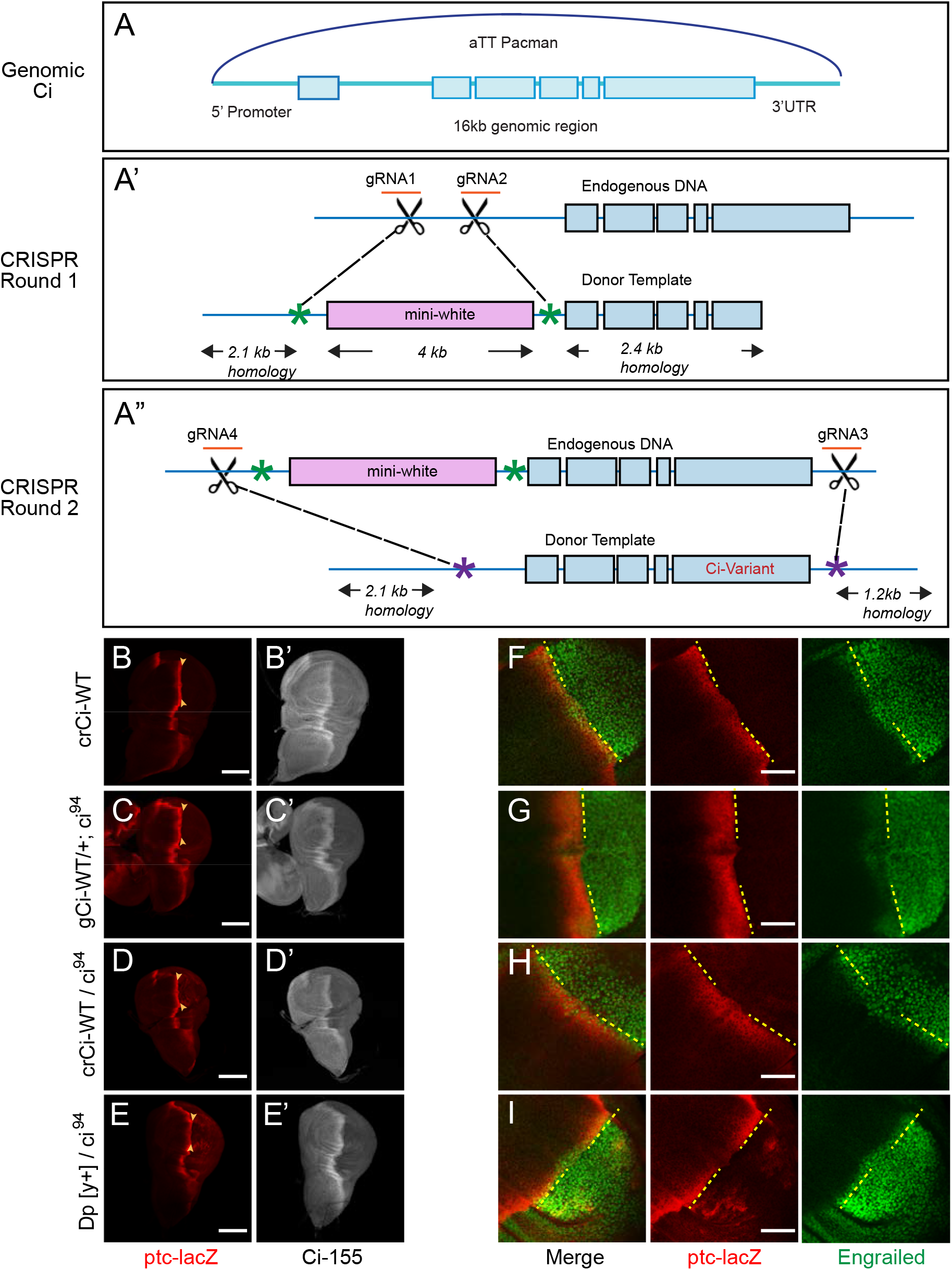
Wild-type genomic ci transgene and CRISPR ci alleles are fully functional. (A) The “genomic Ci” transgene (*gCi*) was derived from a 16kb genomic region of *ci* cloned into an att-Pacman vector and inserted at *att ZH-86FB* located at 86F on the third chromosome. It includes a 6.7kb upstream promoter region, exons1-6 (blue boxes), introns (blue line), and 0.7kb of the 3’UTR. (A’-A”) *CRISPR ci* (*crCi*) alleles were generated in two rounds. (A’) The first round inserted a mini-white gene (pink box) into the first intron of endogenous *ci* using two guide RNAs (orange) in the first intron and altered PAM sites on the donor template (green star). (A”) The second round replaced intron 1 and exons 2-6 (blue lines and boxes) including the mini-white gene; the donor template had mutated PAM sites (purple stars) corresponding to the gRNA4 site, approximately 30 bp outside the mutated PAM site for gRNA1, and in the 3’UTR 5kb away from gRNA 2, labeled gRNA 3. (B-E) Third instar wing discs showing *ptc-lacZ* reporter gene expression, visualized by Beta-galactosidase antibody staining (red), with the posterior edge of AP border expression marked by yellow arrowheads, and (B’-E’) full-length Ci-155, visualized by 2A1 antibody staining (gray-scale). Anterior is left and ventral is up. (B) Two copies and (D) one copy of *crCi-WT*, or (C) one copy of *gCi-WT* supported normal patterns of elevated *ptc-lacZ* and Ci-155 at the AP border but (C, D) sporadic ectopic posterior *ptc-lacZ* expression was seen whenever a single *ci^94^* allele was present, even (E) in discs with no synthetic *ci* transgene or allele (*Dp[y^+^]* has wild-type *ci*). (F-I) Induction of En (green) at the AP border was detected by using the posterior boundary (yellow dashed line) of *ptc-lacZ* (red) to distinguish anterior (left) from posterior compartment cells, which express En independent of Hh signaling. En induction was normal in the presence of (F) two copies of *cr-Ci-WT*, (H) one copy of *crCi-WT* or (I) one wild-type *ci* allele and (G) was slightly reduced in the presence of one copy of *gCi-WT*. Scale bars are (B-E) 100μm and (F-I) 40μm.

We also created mutant *ci* alleles by using CRISPR in two rounds: in the first round, we put a *mini-white* marker gene in the first intron of *ci* (Fig. 1B); in the second round we selected against the *mini-white* gene and introduced our mutation of interest, replacing the DNA between the first intron and the 3’UTR (Fig. 1C). A single copy of *crCi-WT* in combination with *ci^94^* resulted in efficient development of normal adults and larval wing discs with normal patterns of En, *ptc-lacZ* and Ci-155 expression in the anterior compartment (Fig. 1D, F). During these studies we also became aware of a *ptc-lacZ* artifact, whereby low levels of *lacZ* product were detected sporadically in posterior cells of wing discs when there was a single *ci^94^*allele; this occurred in flies with a normal *ci* allele (on the *Dp(y^+^)* “balancer”) (Fig.1G) or with the *crCi-WT* allele (Fig.1F). The artifact was also seen with *gCi-WT* when *ci^94^* was heterozygous (data not shown) but not when *ci^94^* was homozygous (Fig.1E) or when *crCi-WT* was homozygous (Fig. 1D). We sequenced the relevant region of the *ci^94^* allele and confirmed that it was the same deletion originally reported (Methot and Basler, 1999) and in FlyBase. We also induced homozygous *ci^94^* clones (by *FRT*-mediated recombination to remove a second chromosome genomic *ci* transgene) and confirmed that *ci^94^* encoded no detectable Ci-155 protein (data not shown).

### Processing-resistant Ci variants have stabilized Ci-155 levels

PKA phosphorylates Ci-155 at amino acids S838, S856, and S892 to create recognition sites for both GSK3 and CK1, which further phosphorylate Ci-155 at a consecutive series of primed phosphorylation sites (Smelkinson and Kalderon, 2006; Smelkinson et al., 2007). The phosphorylation series creates a binding site for Slimb that consists of the core peptide pSpTYYGpS_849_MQpS spanning residues 844-852. Ci-S849A lacks the last CK1 target site initially primed by PKA phosphorylation of S838 and Ci-P(1-3)A has alterations to all three PKA sites (S838A, S856A, and S892A). Ci fragments with those alterations showed complete loss of Slimb binding in vitro after exposure to PKA, CK1 and GSK3, while *UAS-Ci* transgene products with those changes showed no processing in wing discs, judged by a sensitive assay of repressor function in posterior compartment wing disc cells (Smelkinson and Kalderon, 2006; Smelkinson et al., 2007). We used *ci* alleles with those alterations to determine how loss of processing affects Ci protein levels and activity at normal physiological levels. The wing discs of animals with *crCi-S849A* or *crCi-P(1-3)A* in combination with *ci^94^* had expanded anterior regions (Fig. 2A-D), as expected because Ci-75 repressor, normally produced from Ci-155 processing, is required to silence *dpp* expression in anterior cells and ectopic Dpp induces anterior growth and pattern duplications (Methot and Basler, 1999). We also found that these discs had strongly elevated Ci-155 levels throughout the anterior, indicating that full length Ci-155 was not being processed in the absence of the Hh signal (Fig. 2A-D; Fig. S1A-D, G, H).

**Figure 2.**
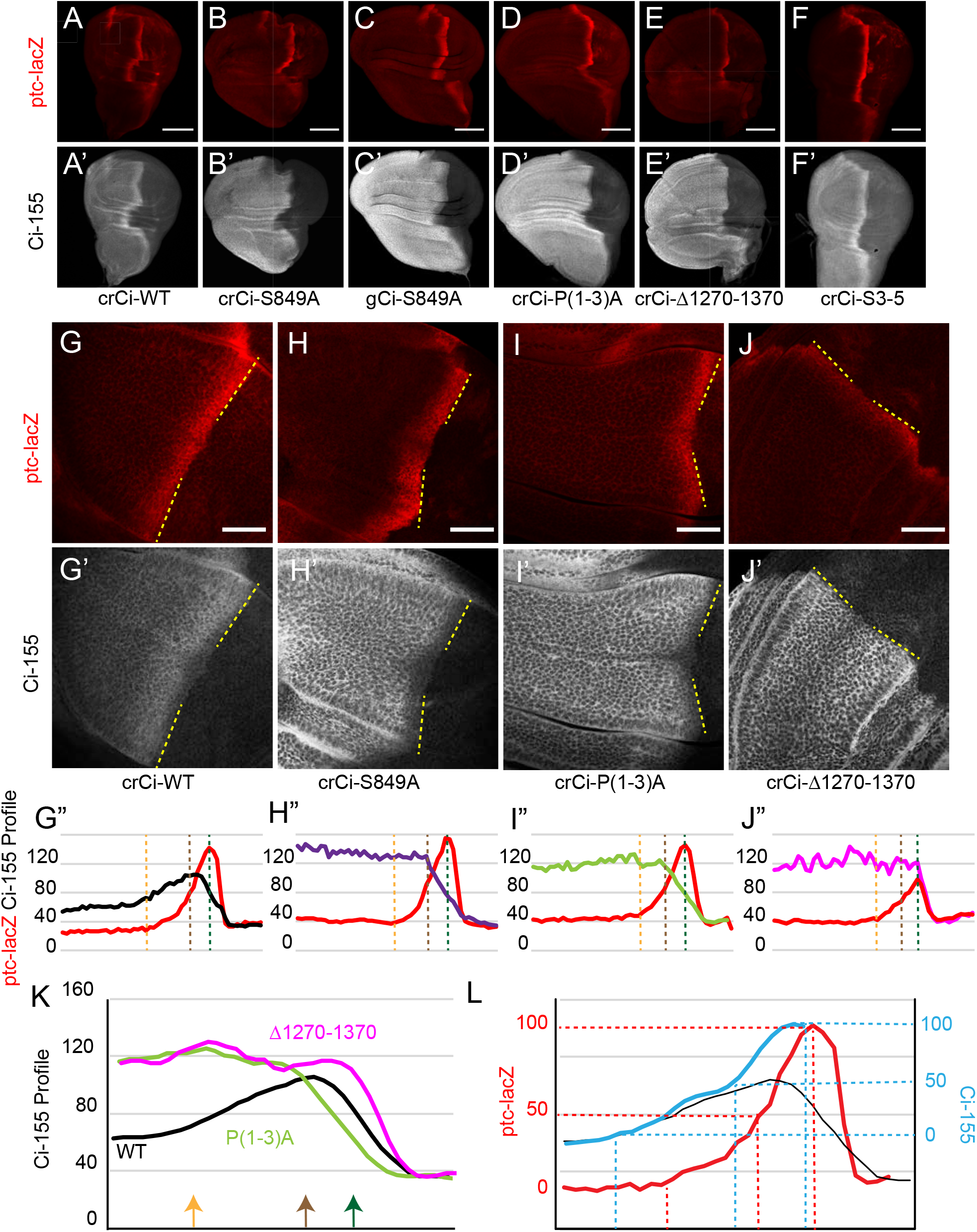
Processing-resistant Ci variants reveal the gradients of Hh-stimulated Ci-155 degradation and Hh inhibition of Ci-155 processing. (A-J) *ptc-lacZ* (red) and (A’-J’) Ci-155 (gray-scale) in wing discs with one copy of the indicated *crCi* alleles (with *ci^94^*) at (A-F) low (20x objective) and (G-J) high (63x objective) magnification, with AP boundary (dotted yellow line at posterior *ptc-lacZ* boundary). Scale bars are (A-F) 100μm and (G-J) 40μm. (G”-J”) Intensity profiles for *ptc-lacZ* (red) and Ci-155 (black) from anterior (left) to posterior. Vertical lines indicate *ptc-lacZ* peak (green), initial rise (orange) and 50% increase to peak (rbrown). (K) Normalized Ci-155 profiles for indicated *crCi* alleles derived from G”-J” but with a smoothing function that calculates average intensity for five successive locations centered on each x-axis location. Arrows indicated locations of *ptc-lacZ* initial rise, 50% increase and peak for *crCi-WT* discs. (L) *ptc-lacZ* (red) and Ci-155 (black) smoothened profiles for *crCi-WT*, with red guide lines for locations of initial rise, 50% increase and peak *ptc-lacZ*. The difference between the average of Ci-155 intensity for Ci-P(1-3)A Ci-S849A at each point along the x-axis was subtracted from the maximum intensity for those genotypes to calculate a value for Hh-stimulated degradation, which was added to the Ci-WT Ci-155 profile at each location to produce the blue curve, representing Ci-155 levels in the absence of Hh-stimulated degradation. Blue guide lines show the locations where inferred Ci-155 processing is first inhibited, 50% inhibited, and fully inhibited.

### Processing-resistant Ci variants reveal Hh-stimulated Ci degradation

Although normal wing discs have a clear stripe of elevated Ci-155 at the AP border relative to anterior cells, Ci-155 levels actually decline over the posterior half of the AP border (Fig. 2G) in a manner that depends on strong activation of the Hh pathway (manifest most clearly by the absence of this effect in wing discs lacking Fu kinase activity) (Ohlmeyer and Kalderon, 1998). Hh-stimulated Ci-155 degradation close to the Hh source has been attributed, at least in part, to induction of *hib*, also known as *roadkill (rdx,)* in a narrow stripe within the strong *ptc-lacZ* domain (Kent et al., 2006; Zhang et al., 2006). Hib is a BTB protein (Broad Complex, Tramtrack, and Bric a Brac) and is the substrate recognition component of a Cul3 E3 ubiquitin ligase complex that promotes full degradation of Ci-155 following ubiquitination. That proteolysis mechanism differs from Ci-155 partial proteolysis (processing) initiated by Slimb within a Cul1 E3 ubiquitin ligase complex (Jiang, 2006).

Although Ci-155 protein levels were strongly elevated compared to normal for Ci-S849A and Ci-P(1-3)A in anterior cells, there was a sharp decline towards the posterior of AP border territory (Fig. 2H, I; Fig. S1A-D, G, H). This profile presumably represents the gradient of Hh-stimulated Ci-155 degradation. It has not previously been seen in isolation because it is super-imposed on a gradient of opposite polarity due to inhibition of Ci processing for wild-type Ci (Fig. 2K). Indeed, subtraction of the processing-resistant Ci-155 profile from the normal Ci-155 profile revealed an inferred gradient of normal Ci-155 processing (Fig. 2L). The deduced profile is similar to that of *ptc-lacZ* activation, with a slightly higher sensitivity to low levels of Hh (Fig. 2L). Thus, comparison of the Ci-155 profiles of wild-type and processing-resistant variants provided the best evidence to date of the spatial profiles of graded inhibition by Hh of Ci-155 processing (Fig. 2L) and of graded Hh-promoted Ci-155 degradation at the AP border (Fig. 2H, I, K).

We also created a *ci* allele resembling a C-terminal deletion variant that had previously been found not to undergo processing in assays using cultured cells and *UAS-Ci* transgenes in wing discs (Wang and Price, 2008; Zhou and Kalderon, 2010). CiΔ1270-1370 also had uniformly elevated Ci levels in anterior wing disc cells, consistent with a lack of processing (Fig. 2E, J; Fig. S1E). However, unlike Ci-S849A and Ci-P(1-3)A, this Ci variant induced *ptc-lacZ* only weakly at the AP border (Fig. 2A-E, G-J; Fig. S1P). There was also no decline of Ci-155 protein within AP territory (Fig. 2J, K), consistent with prior evidence that Hh-stimulated Ci-155 degradation, visualized clearly with the other processing-resistant Ci variants, is only observed at high levels of Hh signaling.

To study Hh-promoted degradation further we created an allele encoding a Ci variant with compromised Hib binding. Hib binds to Ci-155 through multiple sites; altering three principal binding regions (S3,4,5) through clustered point mutations rendered the altered Ci-155 (“Ci-S3-5”) largely insensitive to Hib in a tissue culture assay (Zhang et al., 2009). We found that animals expressing one copy of *crCi-S3-5* (in combination with *ci^94^*) developed efficiently intro adults with normally patterned wings (data not shown). In larval wing discs, *ptc-lacZ* expression was slightly elevated at the AP border peak compared to normal but the domain of induction of En (a high-level Hh target) was not expanded (Fig. 2F; Fig. S1F, I, M-O). The Ci-155 profile included low anterior levels, suggesting normal processing, and declined in the posterior regions of the AP border much like wild-type Ci-155, showing that Hh-stimulated Ci-155 degradation remained robust (Fig. 2F; Fig. S1I, L). It seems likely from the properties of Ci-S3-5 that direct action of Hib on Ci-155 does not account for a significant fraction of the degradation of Ci-155 stimulated by the highest levels of Hh signaling, consistent with results from another study (Seong et al., 2010; Seong and Ishii, 2013).

Suppressor of fused (Su(fu)) has been implicated in Hh-stimulated Ci-155 degradation because Hib can indirectly affect Su(fu) protein levels, loss of Su(fu) leads to greatly reduced Ci-155 levels throughout the wing disc and it has been conjectured that Hh may activate Ci-155 in part through Su(fu) dissociation from Ci-155 (Humke et al., 2010; Liu et al., 2014; Ohlmeyer and Kalderon, 1998; Seong and Ishii, 2013). We examined Ci-155 AP border profiles for wild-type Ci and Ci-S3-5 in the absence of Su(fu). The profiles were extremely similar and Ci-155 levels appeared to peak at, or very close to the AP compartment boundary, suggesting little or no Hh-stimulated degradation (Fig. S1J-L). The results are consistent with the idea that Su(fu) is a key factor in the regulation of Hh-stimulated Ci-155 proteolysis.

### Processing-resistant Ci variants have normal activity

The pattern of En and *ptc-lacZ* induction at the AP border was normal for Ci-S849A and Ci-P(1-3)A (Fig. 3A-C; Fig. S1P), showing that Ci-155 processing is not essential for dose-dependent induction of Hh target genes, However, the expanded anterior regions of these wing discs are likely responsible for the failure to recover adults expressing only processing-resistant Ci variants. These wing discs expressed *dpp* ectopically in anterior cells, as expected, and the presence of a *ci^Ce^* allele, which encodes a constitutive repressor form of Ci (and no activator) (Methot and Basler, 1999) restored normal wing disc morphology and allowed recovery of adults (Fig. S2A-D).

**Figure 3.**
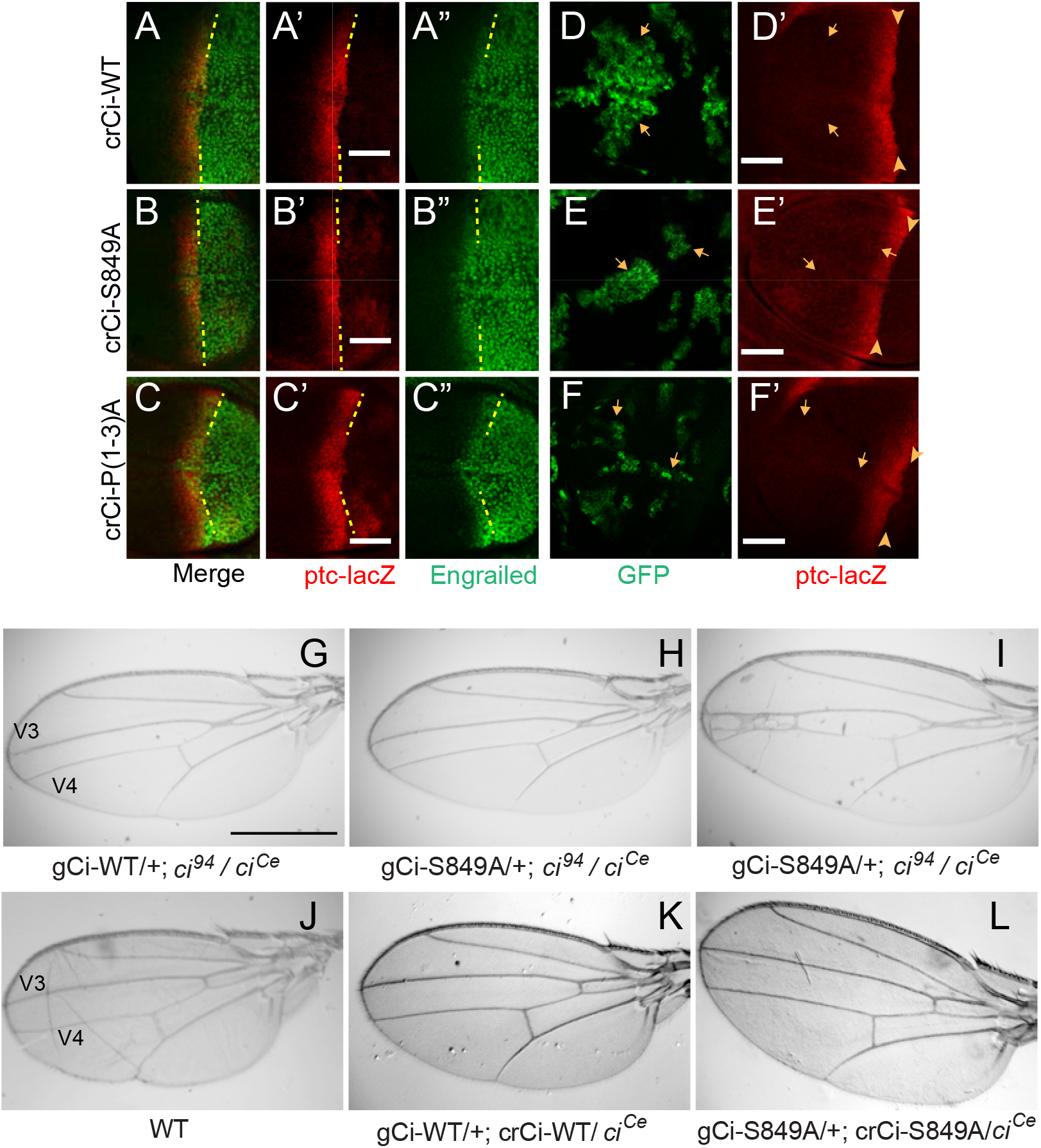
Processing-resistant Ci variants support normal Hh signaling and wing patterning. (A-C) Wing discs with one copy of indicated *crCi* alleles. *ptc-lacZ* (red) indicates the AP compartment boundary (yellow line) to reveal induction of the high-level Hh target gene En (green) in anterior cells at the AP border. (D-F) Wing discs with anterior clones (GFP, green, yellow arrows) that have lost a second chromosome *gCi* transgene, leaving one copy of the indicated *crCi* alleles as a source of Ci. (D’-F’) Little (E’) or no (D’, F’) *ptc-lacZ* induction was observed in the clones (arrows) relative to the AP border (arrowheads). Scale bars are (A-F) 40μm. (G-L) Wings from adult flies with the indicated *ci* transgenes and alleles (*ci^Ce^* encodes a constitutive repressor). The spacing between veins 3 and 4 is (J-L) normal for two copies of WT or S849A *ci* alleles and (G-I) similarly reduced for one copy of WT or S849A alleles. Scale bars are (G-I) 500μm.

Normal wing morphology depends on long-range patterning elicited by the central stripe of Hh-induced Dpp and on creation of a central inter-vein region between veins 3 and 4 by stronger Hh signaling, sufficient to induce the transcription factor Collier (Vervoort, 2000; Vervoort et al., 1999). The adult wing phenotypes of animals with *gCi-WT* or *gCi-S849A* in a *ci^94^*/*ci^Ce^* background were similar to each other, with a consistent moderate pinching between veins 3 and 4 (Fig. 3G, H), though some animals with *gCi-S849A* also showed a greater narrowing of the inter-vein region (Fig. 3I). We then added a *crCi* allele in place of *ci^94^* and found that wing morphology was absolutely normal for flies with either wild-type Ci or Ci-S849A encoded by a *gCi* transgene together with a *crCi* allele in trans to *ci^Ce^* (Fig. 3J-L). Hence, we conclude that Hh can fulfill its normal morphogenetic function, culminating in a normally patterned wing in the complete absence of regulated Ci-155 processing.

### Dependence of Ci-155 activation by Fused kinase on inhibition of Ci-155 processing

At the AP border, Ci-155 processing is inhibited and both Ci-155 and Fused kinase are activated. Genetic elimination of Fu kinase activity prevents En induction, greatly reduces *ptc-lacZ* induction and leads to the emergence of adults with no space between veins 3 and 4 (Ohlmeyer and Kalderon, 1998; Preat et al., 1993). Fu can be activated synthetically in the absence of Hh stimulation by overexpression of Fu variants with either a membrane targeting tag (GAP-Fu) or acidic residue replacements of phosphorylation sites key to normal activation (Fu-EE) (Claret et al., 2007; Zhou and Kalderon, 2011). Activated Fu can also partially activate Smo (Claret et al., 2007; Sanial et al., 2017) but direct downstream, Smo-independent actions can be measured by assaying responses in *smo* mutant anterior clones expressing Fu-EE or GAP-Fu. Previously, such experiments showed that activated Fu alone was sufficient to elicit strong Hh target gene induction, suggesting that Ci-155 activation can be effective even without the normal inhibition of processing that occurs at the AP border (Zhou and Kalderon, 2011). It was also observed, however, that Ci-155 levels were increased by activated Fu, suggesting partial inhibition of Ci-155 processing even though Fu kinase activity is not normally required for processing inhibition at the AP border (Zhou and Kalderon, 2011).

To clarify the dependence of Ci-155 activation by Fu on Ci-155 processing inhibition we compared the activities of wild-type and processing-resistant Ci variants in *smo* mutant clones expressing activated GAP-Fu. We found that Ci-S849A or CiΔ1270-1370, provided by a single *crCi* allele, mediated *ptc-lacZ* induction in anterior *smo GAP-Fu* clones to the same level as at the AP border, whereas induction mediated by wild-type Ci was much lower (about 50%) (Fig. 4A-D, K). Moreover, when slightly lower levels of protein were provided by a single *gCi* transgene *ptc-lacZ* induction was still lower. In each case, the activity in clones was compared to the AP border of the same wing discs and reflects the activity of the same source of Ci. Similar results were seen in GAP-FU clones that retained one functional *smo* allele, with *ptc-lacZ* induction of four processing-resistant variants (Ci-S849A, Ci-P(1-3)A, CiΔ1270-1370 and CiΔ1270-1370) greatly exceeding that of wild-type Ci (Fig. 4F-J, L). Thus, the activity of Ci-155 stimulated by Fu kinase alone was substantially increased if Ci-155 processing was also inhibited. The comparison of the responses of *gCi-WT* and *crCi-WT* (Fig. 4K) also clarifies that more Ci-155 is the key factor, rather than less Ci-75. Thus, producing and preserving a robust supply of Ci-155 is important for Ci-155 activation by Fu.

**Figure 4.**
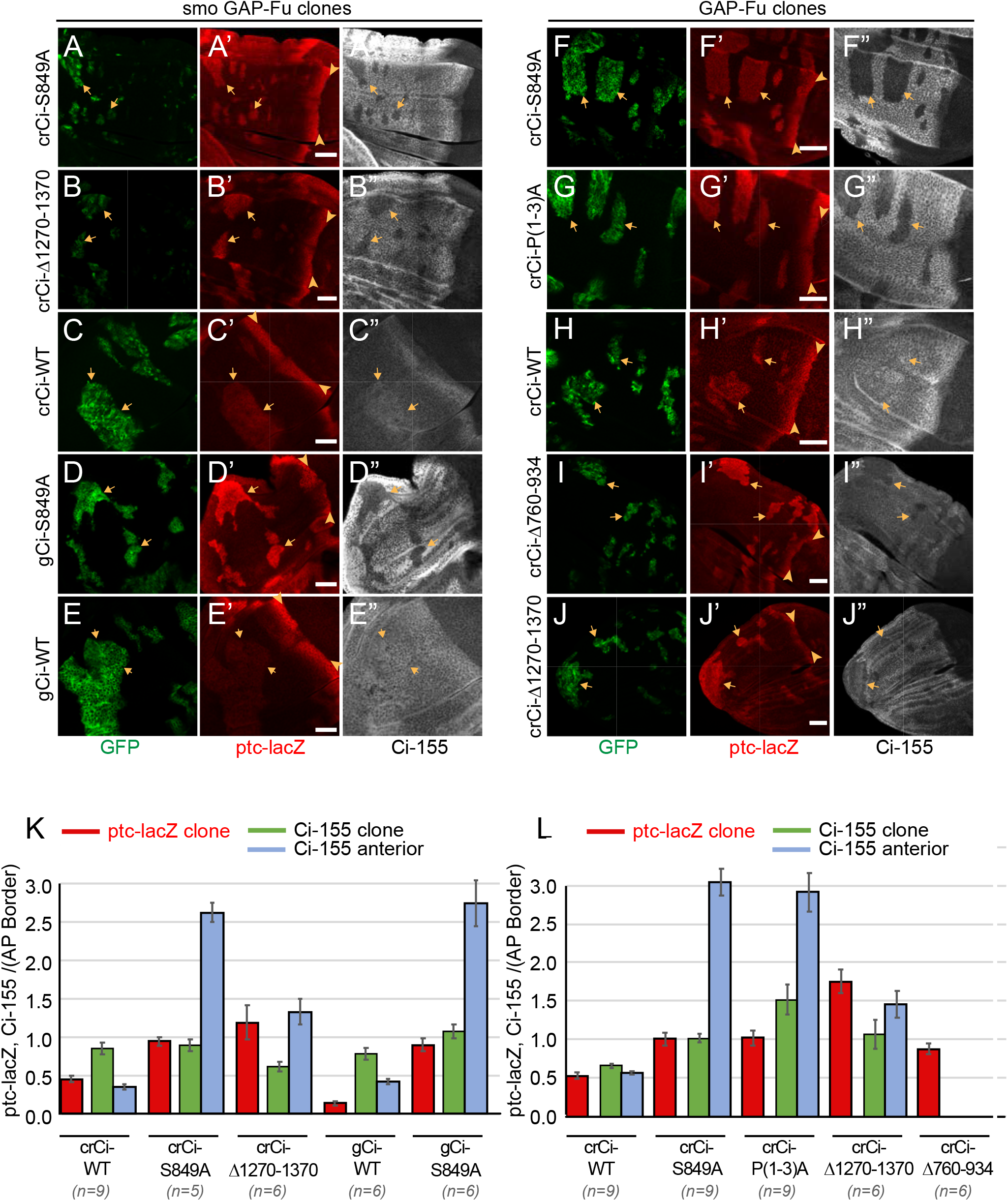
Activation by Fused kinase is enhanced by blocking Ci-155 processing. (A-J) Wing discs from animals with one copy of the designated *ci* transgenes and alleles (together with *ci^94^*) with clones (GFP, green, arrows) that express *UAS-GAP-Fu* and (A-E) lack *smo* activity or (F-J) are heterozygous for *smo* (arrowheads indicate AP border), showing (A’-J’) *ptc-lacZ* (red) and (A”-J”) Ci-155 (gray-scale). (A”-J”) Ci-155 levels were much reduced in clones whenever pathway activity was strongly induced (A”, B”, D”, F”, G”). Scale bars are 40μm. (K, L) Average intensity of *ptc-lacZ* in clones (red), Ci-155 in clones (green) or neighboring anterior territory (blue), as a fraction of AP border levels for (K) *smo GAP-Fu* clones and (L) *GAP-Fu* clones. Mean and SEM shown.

The Ci-155 levels detected within *smo GAP-Fu* clones were much lower than in surrounding territory in wing discs expressing processing-resistant Ci variants (Fig. 4A, B, D, K). The reduction in Ci-155 is similar in magnitude to that observed in posterior regions of the AP border and presumably reflects the Ci-155 degradation normally observed in response to strong Hh stimulation. Wild-type Ci-155 levels were slightly elevated in *smo GAP-Fu* clones relative to neighboring cells but not to the same level as at the AP border (Fig. 4C, K), suggesting possible processing reduction of a magnitude much lower than at the AP border. It is likely there is little or no Ci-155 degradation in these clones because pathway activity is only moderate. Thus, activation of C-155 by Fu to produce high levels of Hh target gene expression requires supplementing Fu actions by providing high levels of primary Ci-155 translation product, protected from processing. The elevated Ci-155 supply produced by processing-resistant Ci variants is, however, not directly evident from measurement of steady-state Ci-155 levels because of subsequent, robust degradation.

### PKA and Cos2 silence Ci-155 activity

It was previously appreciated that Cos2, PKA and Slimb are all necessary for Ci-155 processing but that induction of Hh target genes was higher in anterior *cos2* and *pka* mutant clones than in *slimb* mutant clones (Smelkinson et al., 2007; Wang et al., 1999). The hypothesis that PKA and Cos2 additionally inhibit Ci-155 activity (in the absence of Hh) was supported by the observations that loss of PKA increased *ptc-lacZ* induction in *slimb* mutant clones but it remained possible that there could be some residual processing in the hypomorphic *slimb* clones tested (Smelkinson et al., 2007). Similar tests were conducted using processing-resistant *UAS-Ci* transgenes, also suggesting PKA affects Ci-155 activity, but in both cases the transgene expression level was not physiological and in one case the Ci variant used was still subject to PKA-dependent degradation rather than processing (Smelkinson et al., 2007; Wang et al., 1999).

We therefore induced *pka* and *cos2* clones in wing discs expressing Ci-P(1-3A) from a single allele in combination with *ci^94^*. We found that in both sets of clones, there was a marked increase in *ptc-lacZ* compared to surrounding tissue (Fig. 5B, C, I, L) and compared to clones with no change in PKA or Cos2 activities (Fig. 3F). The level of *ptc-lacZ* induced was found to be about 75% (*pka* clones) or 50% (*cos2* clones) of AP border levels in the same wing discs and in each case matched the levels induced in animals expressing one allele of wild-type Ci (Fig. 5A, H, L). Induction of *ptc-lacZ* was substantially lower for *pka* mutant clones expressing wild-type Ci from only one *gCi* allele (Fig. 5F, L), showing that Ci-155 activation due to loss of PKA activity depends on Ci-155 levels, as previously surmised (Ohlmeyer and Kalderon, 1998) and analogous to the dependence of activation by Fu on processing inhibition. The results clearly indicate that PKA and Cos2 inhibit the activity of Ci-155 that is not processed in the absence of Hh stimulation. The magnitude of inhibition is substantial.

**Figure 5.**
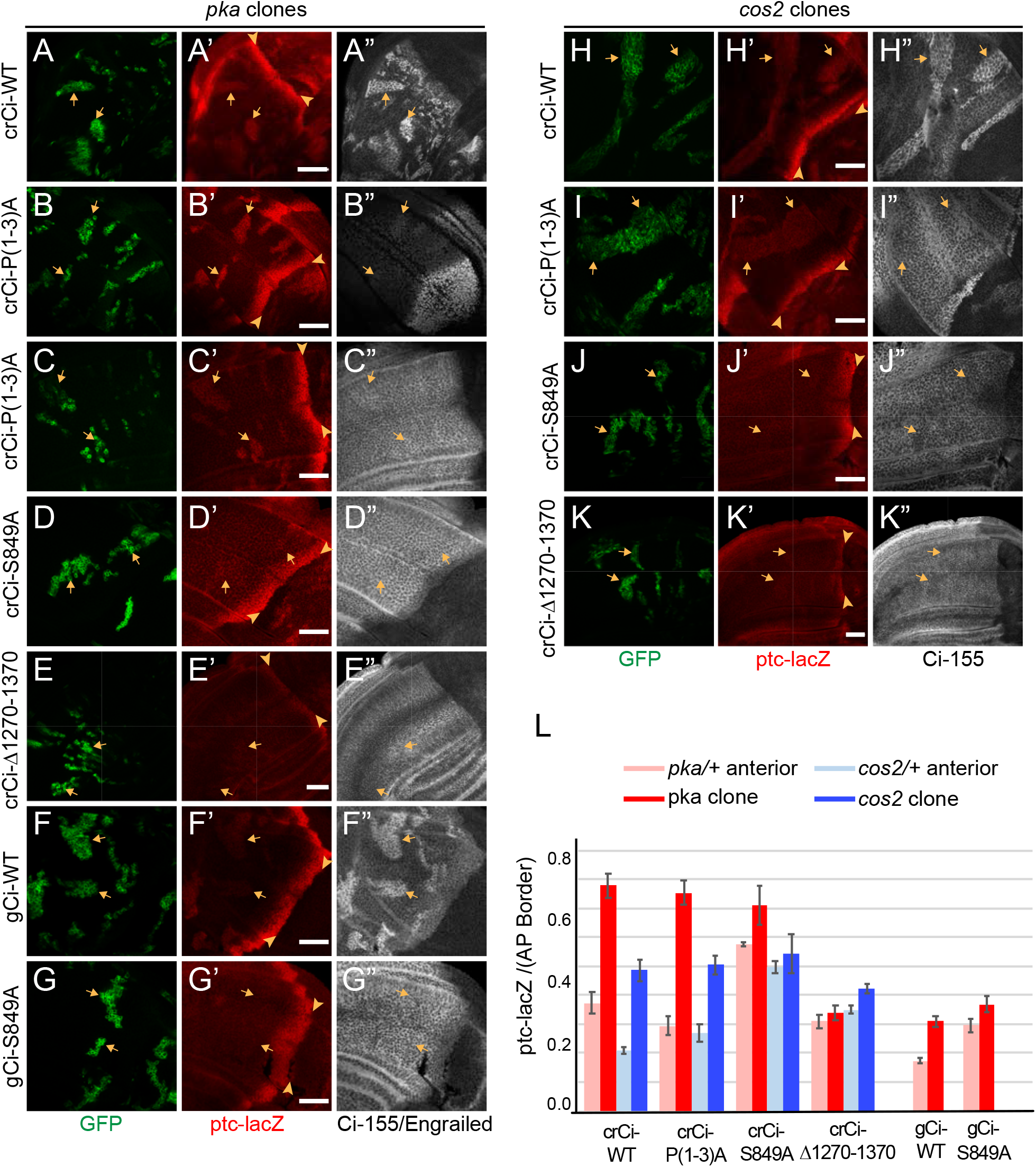
PKA and Cos2 reduce the activity of Ci-155 that is not processed. (A-K) Wing discs from animals with one copy of the designated *ci* transgenes and alleles (together with *ci^94^*) with clones (GFP, green, arrows) that lack (A-G) *pka* activity or (H-K) *cos2* activity (arrowheads indicate AP border), showing (A’-K’) *ptc-lacZ* (red) and (A”, C”-K”) Ci-155 (gray-scale). (B”) En (gray-scale) was weakly induced in *pka* clones from discs expressing Ci-P(1-3)A. Ci-155 levels were (A”, F”, H”) increased relative to neighboring anterior territory for Ci-WT but were (C”-E”, G”, I”-K”) either unchanged or slightly reduced, presumably from full proteolysis, for processing-resistant Ci variants. Scale bars are 40μm. (L) Average intensity of *ptc-lacZ* in *pka* clones (red) or neighboring anterior territory (pink), and in *cos2* clones (dark blue) or neighboring anterior territory (light blue), as a fraction of AP border levels. Mean and SEM shown.

Surprisingly, Ci-S849A activity was not greatly activated by loss of PKA or Cos2 and there was also some *ptc-lacZ* expression in heterozygous tissue surrounding clones (Fig. 5D, G, J, L), similar to the low level of *ptc-lacZ* activity observed for Ci-S849A in clones with normal PKA and Cos2 activity (Fig. 3D, E), suggesting that S849 is relevant to the regulation of Ci-155 activity by PKA and Cos2. CiΔ1270-1370, which induces lower levels of *ptc-lacZ* at the AP border than Ci-WT but is strongly activated by GAP-Fu, was also not activated by loss of PKA or Cos2 (Fig. 5E, K, L). We speculate that perhaps CiΔ1270-1370 is deficient for all responses other than strong stimulation by Fu because it is inhibited more strongly than wild-type Ci by Su(fu).

### Cos2 silences Ci activity by binding to the CORD region

To investigate how Cos2 can silence Ci-155 activity, we further examined the interactions between these proteins. Cos2 can bind to Ci-155 through three regions defined by in vitro binding assays: the CDN region (residues 346-440), the zinc fingers (residues 506-620) and the CORD domain (residues 934-1065) (Wang and Jiang, 2004; Zhou and Kalderon, 2010). Measurement of processing through Ci-155 levels and generation of repressor activity from *UAS-Ci* transgenes in wing discs previously showed that processing was unaffected by removal of the CDN or CORD domain alone, or even when both were deleted but was abrogated when the zinc fingers were also deleted, suggesting that any of the three Cos2-binding domains can suffice for efficient Ci-155 processing (Zhou and Kalderon, 2010). To test which of these domains might be responsible for inhibiting Ci-155 activity we generated *ci* alleles lacking CDN, CORD or both regions. No mutations have been identified that abolish Cos2 binding of the zinc finger region without also preventing DNA binding and consequent Ci activity.

We induced *pka* and *cos2* clones in wing discs expressing only CiΔCORD, CiΔCDN or CiΔCORDΔCDN.^.^ We found that *ptc-lacZ* levels in *cos2* mutant clones were very similar in each case to those mediated by wild-type Ci (Fig. 6 G-K). Those results are consistent with the deletions not affecting any Ci-155 activity other than binding to Cos2 (there is no Ci-75 repressor potentially generated in any clones because there is no Ci-155 processing). In *pka* mutant clones the level of *ptc-lacZ* was similar for wild-type Ci and CiΔCDN but it was significantly higher for CiΔCORD and CiΔCORDΔCDN, and it was also elevated for CiΔCORD expressed from a *gCi* transgene (Fig. 6A-F, K). These observations suggest that Cos2 can inhibit Ci activity specifically by binding to the CORD region of Ci-155. They also show that Ci-155 activation by loss of PKA activity and loss of Cos2-CORD binding can be additive.

**Figure 6.**
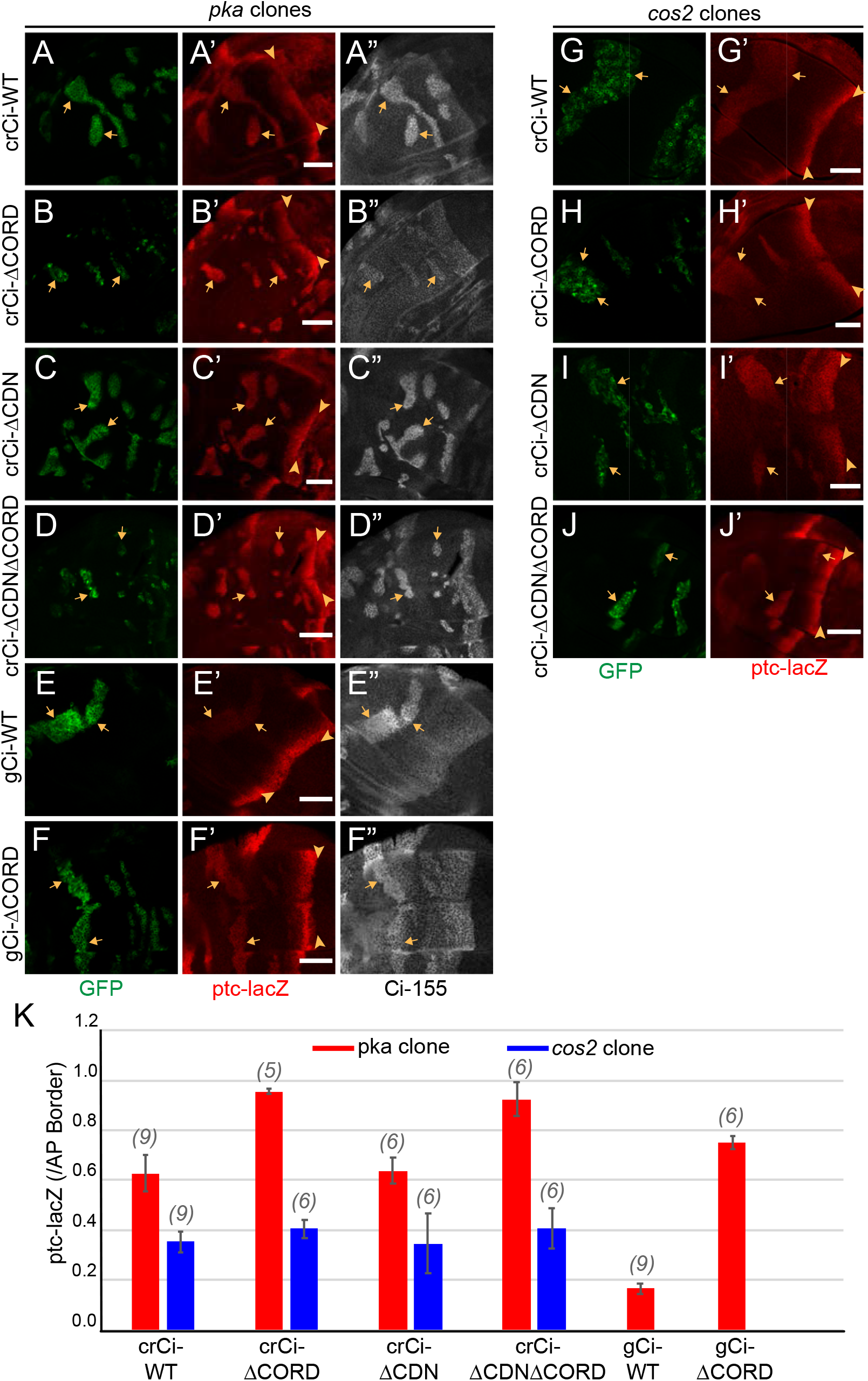
Cos2 reduces Ci-155 activity by binding to the CORD region. (A-J) Wing discs from animals with one copy of the designated *ci* transgenes and alleles (together with *ci^94^*) with clones (GFP, green, arrows) that lack (A-F) *pka* activity or (G-J) *cos2* activity (arrowheads indicate AP border), showing (A’-J’) *ptc-lacZ* (red) and (A”-F”) Ci-155 (gray-scale). (A”-F”) Ci-155 levels were increased relative to neighboring anterior territory for all Ci proteins but the increase was relatively small for (B”) Ci-ΔCORD, suggesting that processing outside the clones may be inefficient. By contrast, a large change was observed for Ci-ΔCDNΔCORD, suggesting very efficient processing. Scale bars are 40μm. (K) Average intensity of *ptc-lacZ* in *pka* clones (red) and in *cos2* clones (blue), as a fraction of AP border levels. Mean and SEM shown.

In wing discs with no mutant clones, CiΔCORD, CiΔCDN and CiΔCORDΔCDN all supported a near-normal Ci-155 profile, indicating substantially normal regulation of Ci-155 processing, and strong *ptc-lacZ* expression confined to the AP border (Fig. 7A-D). The slight enhancement of anterior Ci-155 for CiΔCORD was also seen in a *pka* heterozygous background (Fig. 6B) and a Su(fu) mutant background (Fig. S3A, C). It may indicate a mild processing deficit but there was a very strong contrast between anterior and AP border Ci-155 levels of CiΔCORDΔCDN (Fig. 6D; Fig. 7D), supporting previous evidence that Ci-155 lacking both of these Cos2-binding domains is processed very efficiently, perhaps even more efficiently than wild-type Ci, and that Hh blocks processing efficiently (Zhou and Kalderon, 2010).

**Figure 7.**
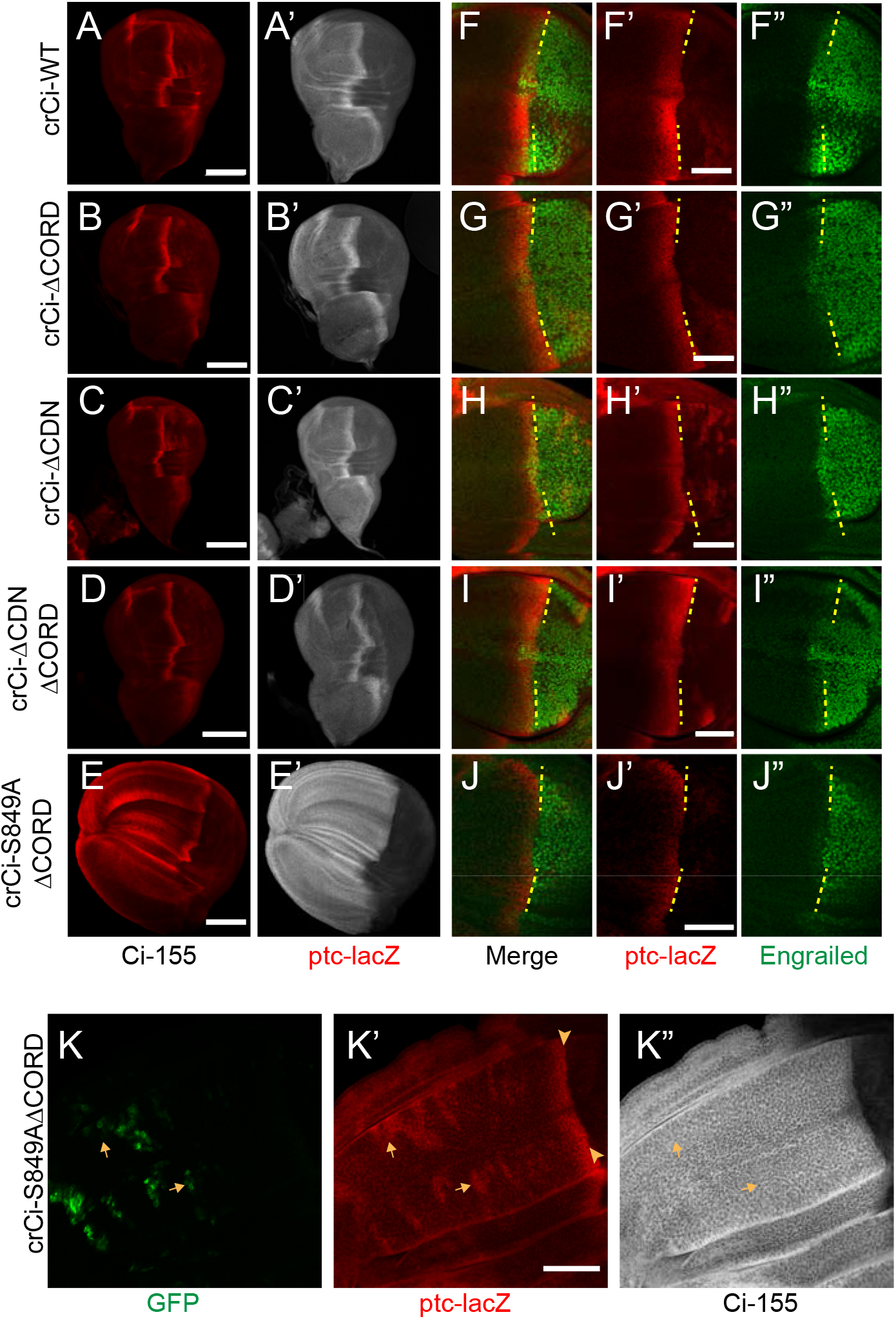
Loss of both Cos2 inhibition and processing combine to activate Ci-155. (A-J) Wing discs from animals with one copy of the designated *ci* alleles (together with *ci^94^*), have (A-D) no ectopic anterior *ptc-lacZ* (red) unless (E) both the CORD domain is removed and processing blocked (by the S849A alteration). (A’-E’) Ci-155 (gray-scale) in the same wing discs. (F-J) Anterior En (green) induction, revealed by marking the AP compartment boundary (yellow lines) with the posterior extent of *ptc-lacZ* (red), was reduced for Ci variants (G, J) lacking the CORD domain, (J) especially together with the S849A alteration. (K) Wing disc with anterior clones (GFP, green, yellow arrows) that have lost a second chromosome *gCi* transgene, leaving one copy of *crCi-S849AΔCORD* as the only source of Ci, showing (K’) *ptc-lacZ* induction in the clones (arrows) to levels similar to the AP border (arrowheads); (K”) Ci-155 (gray-scale) is uniformly high because of blocked processing. Scale bars are (A-J) 40μm.

**Figure 8.**
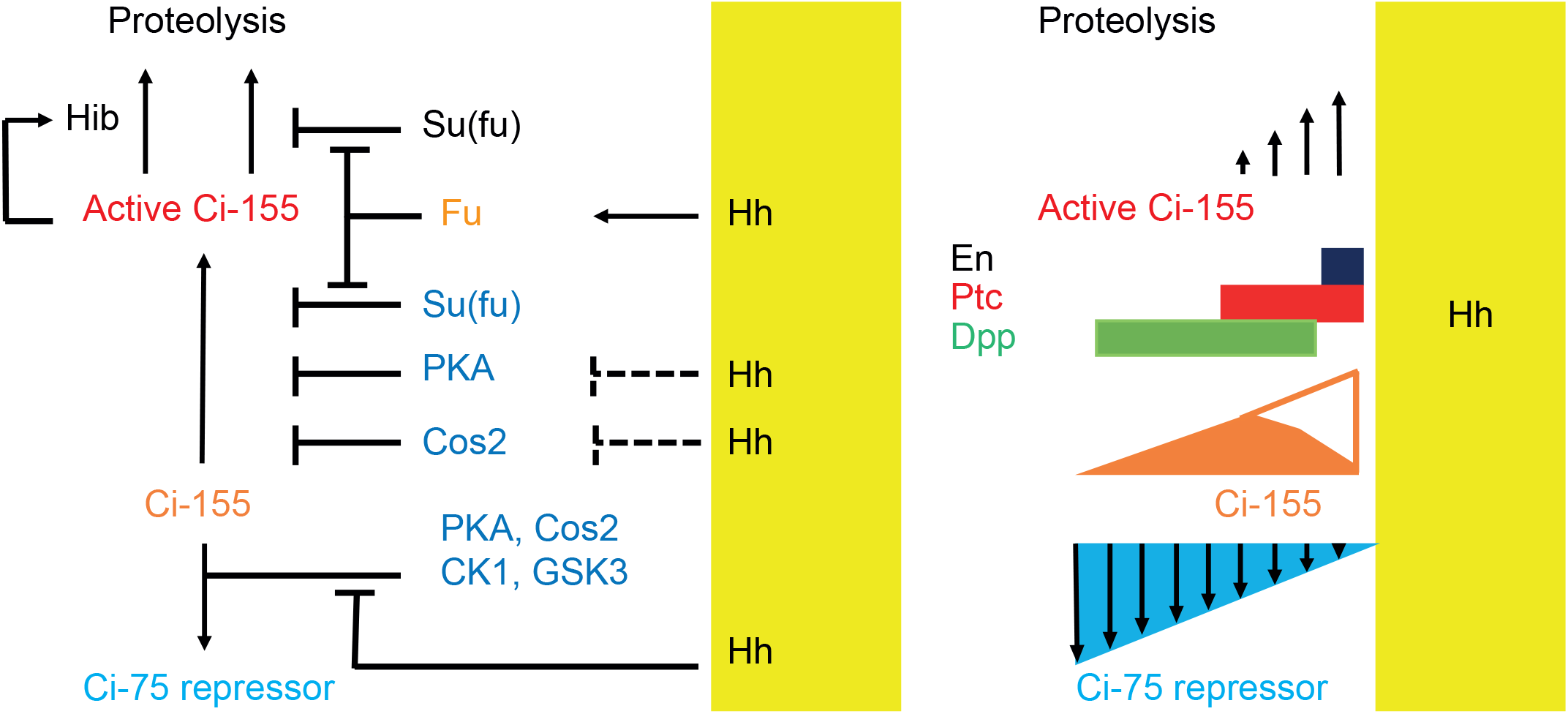
Summary of graded processing, activation and proteolysis of Ci-155 and underlying mechanisms at the AP border. At the AP border, Hh (emanating from posterior, mustard yellow, territory) inhibits Ci-155 processing stimulated by Cos2, PKA, CK1 and GSK3, activates Fu protein kinase activity to activate Ci-155, overcoming inhibition by Su(fu) and other factors, and promotes full Ci-155 proteolysis through induction of Hib and perhaps altering Su(fu) association. Here we used processing-resistant Ci variants to show that PKA and Cos2 (through binding the CORD domain on Ci-155) reduce Ci-155 activation in anterior cells and (right) to deduce the profiles of Hh-stimulated Ci-155 proteolysis (upward arrows) and inhibition of Ci-155 processing (brown and blue triangles) at the AP border that underlie steady-state Ci-155 levels (brown). Graded target gene (En, Ptc, Dpp) activation normally responds to graded levels of activated Ci-155 and Ci-75 repressor but was still observed when there was no regulation of Ci-155 processing and also when the major binding sites for Hib were eliminated, indicating that Ci-155 activation must be graded.

The absence of ectopic anterior *ptc-lacZ* in wing discs expressing CiΔCORD (Fig. 7B, D) suggests that loss of Cos2-CORD association only leads to Ci-155 activity when Ci-155 processing is also inhibited. To test this hypothesis further we created an allele expressing a processing-resistant Ci variant (S849A) that also lacked the CORD domain. We found that, unlike CiΔCORD, Ci-S849AΔCORD in combination with *ci^94^* resulted in wing discs with expanded anterior compartments and ectopic *ptc-lacZ* throughout the anterior (Fig. 7B, E). Ectopic *ptc-lacZ* was much stronger than observed for Ci-S849A and was also evident cell autonomously in clones lacking a wild-type Ci transgene within wing discs expressing Ci-S849AΔCORD (Fig. 7K). Thus, detectable Ci-155 activation by loss of Cos2-CORD interaction requires inhibition of Ci-155 processing.

Animals with the Ci-S849AΔCORD allele were not easily maintained in combination with a wild-type *ci* allele, presumably because Ci-75 activity only just sufficed to prevent significant ectopic Hh target gene activation. Given the high constitutive activity of Ci-S849AΔCORD, we were surprised to find that larval wing discs with this allele and *ci^94^* showed very little induction of En at the AP border (Fig. 7J). A lesser defect was seen for CiΔCORD (Fig. 7A-B) but no such deficit was seen for Ci-S849A (Fig. 3B). To try to understand the origin of this deficit we examined the responses of CiΔCORD to loss of Fu kinase, loss of Su(fu) or both. Loss of Su(fu) did not restore robust En expression (Fig. S3B, D). Induction of *ptc-lacZ* was much reduced by loss of Fu kinase and restored by additional loss of Su(fu), just as observed for Ci-WT (Fig. S3E-H). Wing discs lacking both Fu kinase and Su(fu) did not exhibit AP border En for CiΔCORD or Ci-WT (Fig. S3G, H), consistent with an earlier report that Ci-155 activation by Fu operates substantially, but not entirely by antagonizing inhibition by Su(fu) (Zhou and Kalderon, 2011). We also found that CiΔCORD responded to activated GAP-Fu but did not induce *ptc-lacZ* to higher levels than Ci-WT despite the absence of the potentially inhibitory CORD domain (Fig. S3I-L, N). We speculate that Fu activated by Hh at the AP border may not be brought to activate Ci-155 efficiently in the absence of the CORD domain but excess GAP-Fu can largely compensate for that deficiency. The lesser induction of En by Ci-S849AΔCORD compared to CiΔCORD at the AP border (Fig. 7G, J) suggests an additional speculation that those Ci-155 molecules spared from processing but failing to engage with activated Fu might compete with activated Ci-155 and thereby limit Hh target gene induction.

## Discussion

Hh signaling in Drosophila and in mammals involves two key changes: inhibition of the proteolytic processing of Ci/Gli proteins to repressor forms, thereby also increasing full-length protein levels, and activation of full-length Ci/Gli proteins. The relative importance of repressor and activator, both of which can potentially regulate the same set of genes, varies in different mammalian tissues in part because of the Gli protein expressed (Gli3 is more efficiently converted to repressor than Gli2) and because Gli1 is itself a Hh target gene, acts only as an activator and therefore has a specialized amplification role (Briscoe and Therond, 2013; Kong et al., 2019; Liu, 2019). In Drosophila, Ci is the only transcriptional effector of Hh signaling, allowing straightforward interrogation of the relative importance of regulation through altering the levels of Ci-75 repressor, latent Ci-155 activator and conversion of Ci-155 to a potent transcriptional activator. Moreover, wing disc development is perhaps the most demanding and easily perturbed patterning challenge for Hh signaling in Drosophila and therefore suitable for dissecting essential regulatory influences that support dose-dependent responses. It is therefore perhaps surprising that we found that regulation of Ci-155 processing is not essential for major manifestations of Hh morphogen action in wing discs.

### Evidence that regulation of Ci-155 processing is not essential for Hh morphogen action

We found normal patterns of *ptc-lacZ* and En induction at the AP border supported by a Ci variant that cannot be processed. We tested only one copy of the *ci-S849A* and *ci-P(1-3)A* alleles, so it remains possible that two copies might impair patterning. We also found that one *crCi-S849A* allele together with a genomic *gCi-S849A* transgene and the constitutive repressor allele *ci^Ce^* produced adults with normally patterned wings, showing that normal Hh signaling does not require regulation of the level of either repressor or full-length Ci protein by processing. The result also shows that high-level signaling is possible without eliminating Ci repressor. Normal wing pattern does not depend on En induction in wing discs and we did not examine En induction directly in this genotype, so it is uncertain whether the highest levels of Hh signaling were reproduced.

It has previously been shown that animals lacking both Fu kinase and Su(fu) activities develop normal wings (although En is not induced in those wing discs) (Ohlmeyer and Kalderon, 1998; Preat, 1992), so robust morphogen responses are also possible in the absence of Fu kinase, the most prominent regulator of Ci-155 activation. Thus, while we have found here that regulation of Ci-155 processing is dispensable, it is possible that a similar conclusion applies to regulation of Ci-155 activation, suggesting redundancy rather than pre-eminence of one mechanism over another. The spatial morphogen action of Hh is aided by the negative feedback loop of *ptc* transcriptional induction leading to increased Hh sequestration by Ptc protein (Chen and Struhl, 1996). That feature not only influences the normal Hh signaling gradient but can dampen changes due to an aberrantly sensitive or insensitive transduction system, permitting less or more Hh spread, respectively, and potentially accommodating the loss of one mode of Ci activity regulation.

### Hh-stimulated Ci-155 degradation

Both the mechanism and the purpose of Hh-stimulated Ci-155 degradation remain elusive (Kent et al., 2006; Liu et al., 2014; Seong and Ishii, 2013; Zhang et al., 2006). Here we have produced the clearest images to date of the magnitude and spatial profile of this process by using physiologically expressed processing-deficient Ci variants to eliminate the normally confounding counter-gradient of Ci-155 processing inhibition. The effect on Ci-155 levels is large, so a significant consequence might be anticipated. However, studies of the effects of eliminating or reducing the activity of the best-publicized mediator Hib/Rdx of Ci-155 degradation have shown mixed results, with no strong reproducible effects on limiting the domain or intensity of the strongest Hh responses in wing discs (Kent et al., 2006; Liu et al., 2014; Seong and Ishii, 2013; Zhang et al., 2006). Here we used a Ci variant with multiple alterations to sites of Hib association, which had previously been shown to almost eradicate direct down-regulation of Ci-155 by Hib E3 ligase complexes under synthetic conditions (Zhang et al., 2009). This Ci variant supported a normal pattern of Hh target gene induction in wing discs and the development of adults with normal wings. The *ptc-lacZ* profile observed in larval wing discs suggested a small increase of *ptc* induction close to the source of Hh but the Ci-155 profile appeared unaltered with robust Hh-stimulated Ci-155 degradation in the posterior part of the AP border. Hib may also affect Ci-155 degradation indirectly by altering Su(fu) protein levels (Liu et al., 2014) or in other ways. However, substantial Hh-promoted Ci-155 degradation may be independent of Hib. The complete absence of Su(fu) greatly reduces Ci-155 levels throughout wing discs (Ohlmeyer and Kalderon, 1998) and it is commonly hypothesized that signaling elicits Ci-Su(fu) dissociation, as suggested by studies of mammalian Hh signaling (Humke et al., 2010; Tukachinsky et al., 2010). Such dissociation, stimulated in proportion to Fu activation would promote Ci-155 degradation in proportion to Ci-155 activation without the necessary participation of a transcriptionally induced intermediate, such as Hib. Consistent with this hypothesis, no reduction of Ci-155 levels close to the source of Hh was apparent in wing discs lacking Su(fu) but the sensitivity of those measurements was limited by the low Ci-155 levels throughout such wing discs. Whether Ci-155 activation by Fu does involve dissociation of Su(fu) and whether that contributes significantly to Ci-155 degradation remain to be thoroughly investigated.

### Ci-155 activation by Fu

We found that artificially activated Fu can activate processing-resistant Ci in anterior, Hh-free territory as effectively as normal Fu activity at the AP border but wild-type Ci was comparatively poorly activated, especially if it was produced at slightly lower levels by a *gCi* transgene rather than a normal *ci* allele. The latter result suggests that the level of Ci-155 is critical for a strong response to Fu rather than reduction or elimination of Ci-75 repressor. One possible hypothesis to account for these results is that Fu can only effectively activate Ci-155 that is free from its major binding partners, Su(fu) and Cos2. However, activation by Fu is not significantly affected by the absence of Su(fu) or excess Su(fu) (Zhou and Kalderon, 2011), suggesting that Fu can counter any influence of Su(fu). Also, Cos2 and Fu stability are inter-dependent (Ruel et al., 2003; Zadorozny et al., 2015), suggesting that even activated Fu may remain with Cos2, thereby ensuring the presence of Cos2 wherever Fu acts.

How Fu activates Ci-155 is not yet clear. Although Fu phosphorylates both Su(fu) and Cos2 in response to Hh, the specific phosphorylation sites identified were found to be unimportant for Hh signaling when assayed under physiological conditions (Oh et al., 2015; Raisin et al., 2010; Ruel et al., 2007; Zadorozny et al., 2015; Zhou and Kalderon, 2011), suggesting that Ci-155 itself may be a crucial direct target of Fu kinase. Specific Fu target sites on Ci-155 have recently been identified and found to alter activity under some conditions but their contributions have not yet been assessed under physiological conditions (Han et al., 2019), an important proviso given the crucial role of physiological conditions for re-evaluating the role of phosphorylation sites on Cos2 (Ranieri et al., 2012; Zadorozny et al., 2015). If Ci-155 is a key Fu target it would be expected that Cos2 might promote that interaction rather than hinder it, further arguing against a requirement for Cos2-free Ci-155 as a substrate.

It therefore seems more likely that high Ci-155 levels are important for Ci-155 activation because signaling is dynamic with a relatively high Ci-155 throughput enforced by robust degradation of activated Ci-155. Indeed, the steady-state level of processing-resistant Ci-155 was dramatically reduced by Fu activation. In other words, the high rate of Hh pathway-induced Ci-155 degradation necessitates a constant supply of fresh Ci-155. This imposed requirement might be the major purpose of Hh-promoted degradation, rather than modulating the profile of the Hh signaling gradient. Under this arrangement, cells will continue to express Hh target genes only when constantly stimulated. The arrangement also allows for the possibility of modulating pathway activity through both the degree of Fu activation and the rate of supply of Ci-155.

### Ci-155 activity regulation by PKA and Cos2

We used the processing-deficient variant Ci-P(1-3A) to show that genetic removal of PKA or Cos2 substantially increased Ci-155 activity in the absence of Hh, providing clear evidence that both PKA and Cos2 inhibit Ci-155 activation in addition to their well-established roles of promoting Ci-155 processing. An unresolved issue is to what extent Hh antagonizes these inhibitory influences.

We found that removal of the CORD domain of Ci conferred significantly higher activity on a processing-resistant Ci variant and increased the response of otherwise normal Ci to loss of PKA, but not to loss of Cos2. These observations are consistent with the hypotheses that the CORD domain is the major mediator of the inhibitory action of Cos2 and that PKA inhibits Ci-155 through an independent mechanism (such that relief of inhibition by both PKA and Cos2 is additive). The CDN domain did not appear to be important for inhibition by Cos2. It is curious that Ci-155 processing may be mediated by any one of three Cos2-binding domains (Zhou and Kalderon, 2010), while Ci-155 inactivation by Cos2 may be mediated just by the CORD domain.

When Hh signals, Ci-155 processing is believed to be reduced primarily through partial dissociation of Cos2-Ci complexes (Li et al., 2014; Ranieri et al., 2014) and this processing inhibition occurs even in the absence of Fu kinase activity (Ohlmeyer and Kalderon, 1998; Smelkinson and Kalderon, 2006). It is not clear which Cos2 interaction domains might be affected the most, what degree of dissociation is elicited or whether Fu kinase might potentiate dissociation. Since *ptc-lacZ* levels in *cos2* mutant clones are much higher than at the AP border of wing discs lacking Fu kinase activity, we can surmise that Fu-independent Cos2-CORD dissociation stimulated by Hh must be only partial. Relief of Cos2 inhibition together with PKA inhibition plausibly account for the low levels of *ptc-lacZ* observed in Fu kinase-deficient discs because processing-resistant Ci-P(1-3)A alone in anterior cells did not induce any *ptc-lacZ* (from one allele). It is therefore likely that the substantial increase in Ci-155 activity observed by genetic elimination of Cos2 and PKA is only partially reproduced by antagonism of inhibitory Cos2 and PKA actions by Hh at the AP border.

The fact that loss of PKA increased the activity of Ci-P(1-3A) shows that the PKA sites used to direct processing are not relevant for the observed inhibitory effect of PKA. Ci-155 includes two additional consensus sites at residues 962 and 1006. In earlier studies using multiple *UAS-Ci* transgenes at a variety of genomic locations, Ci variants lacking all five PKA sites (P1-5A) were found to be more active than those lacking just sites P1-3 (Price and Kalderon, 1999). The relative levels of *ci* transgene expression were not measured in that study and all were likely higher than physiological levels. In mouse studies evidence was provided, albeit with non-physiological expression levels, that alteration of PKA sites in Gli2 analogous to residues 962 and 1006 in Ci-155 increased Gli2 activity (Niewiadomski et al., 2014). We were unable to recover a *crCi* allele encoding a variant with all five PKA sites altered. We were similarly unable to recover variants with processing-resistant alterations together with Su(fu) binding site alterations and the processing-resistant variant with a CORD domain deletion was difficult to recover and propagate. These difficulties were all conceivably because of constitutively high activity, providing some hints that PKA sites 962 and 1006 might be important to restrain Ci-155 activity. However, both of these sites are within the CORD domain and Ci lacking the CORD domain is more strongly activated by loss of PKA than by loss of Cos2, indicating that inhibition by PKA can be mediated by a target that is not one of the five consensus PKA sites in Ci-155. There may, of course, be more than one target through which PKA inhibits Ci-155 activation. If Ci-155 were the major direct target for PKA inhibition we could expect that Hh might relieve this inhibition to roughly the same degree as inhibition by Cos2 (because PKA is brought to Ci by Cos2). Since the PKA target has not yet been defined it remains possible that Hh might not alter inhibition by PKA at all.

## Acknowledgments

This work was supported by NIH RO1 GM041815 to DK. We thank Aaron Choi, Jason Li and Sarah Finkelstein for research assistance, Hoyon Kim and other lab members for continued discussions and input, Rebecca Delker and Dr. Richard Mann for advice on CRISPR engineering, the Bloomington stock center for provision of genetic reagents, the Developmental Studies Hybridoma Bank (DSHB) for antibodies, FlyBase as an information resource, and the confocal microscope resource provided by the Dept. of Biological Sciences, Columbia University.

## Materials and Methods

### Genomic Ci Cloning

Genomic transgenes were created by cloning the entire 16kb genomic *ci* region from a Bluescript-SK (BSK) vector (provided by Dr. K. Basler (Methot and Basler, 1999)) into an *att-Pacman* Expression vector (DGRC). To facilitate mutagenesis, the 16kb fragment was first separated into two parts. The region including the promoter, first exon and part of the first intron (“Ci fragment 2”) was cloned as a BamHI-NheI fragment into BSK cut with BamHI and XbaI to create BSK-CiF2. The complementary NheI-KpnI fragment containing all other exons and the 3’ UTR (“Ci Fragment 1”) was cloned into BSK cut with SpeI and KpnI to create BSK-CiF1. BSK-CiF2 was cut with NotI and Bsp1201 to clone the whole CiF2 fragment into the P[acman]-CmR vector cut with NotI, so that RsrII and PmeI vector sites were downstream of *ci* first intron sequences in RP-CiF2. CiF1 was amplified from BSK-CiF1 by long-range PCR using PfuUltraII Fusion HS DNA polymerase (Agilent Technologies), adding RsrII and PmeI at either end and cloning the product into a Zero Blunt Topo cloning vector (Invitrogen). The RsrII-PmeI fragment was then cloned into RP-CiF2 cut with the same enzyme to create the final Pacman vector containing the entire 16kb genomic ci DNA. The 28kb *gCi attPacman* transgene was then inserted at the *att ZH-86Fb* landing site at cytological location 86F8 (Rainbow Transgenic Services).

### Cloning for generating CRISPR alleles

#### 1^st^ round of CRISPR

A 5kb *mini-white* gene from the *attPacman* construct was cloned into the first intron of “Ci Fragment 1” with the enzyme AaII. The PAM sites associated with guide RNA 1 (TGG->TGA) and guide RNA 2(TGG->TTG) were mutated on “Ci-Fragment 1” in the Ci first intron. guide RNA 1 TCACCCAAAAATCTCGTATT and guide RNA 2 ATATATATACAAGAGTTCCT were cloned in pU6 chiRNA vectors separately. The donor template, guide RNA 1, and guide RNA 2 were then co-injected into fly embryos (*wlig4; attp40 [nos-Cas9]/Cyo*). Flies and guide RNA vectors were obtained from the Mann Lab and injections were carried out using Rainbow Transgenic Services. The injected flies were crossed to *yw hs-flp; Sp/Cyo; TM2/TM6B; Dp[y+]/Dp[y+]* flies (*Dp[y^+^]* is used throughout as an abbreviation for *Dp(1;4)1021[y^+^]*) and progeny screened for male flies that were white^+^. The transformants were balanced and further genotyped to confirm correct placement of the *mini-white* gene (reverse coding orientation compared to *ci*) in the intron. The *ci-[w^+^]* flies (4^th^ chromosome) were used to create a stock, *wlig4; attp40 [nos-Cas9]/Cyo; ci-[w^+^] / ci-[w^+^]*.

#### 2^nd^ round of CRISPR

“Ci Fragment 1” was repurposed as donor construct by adding 500 bp extra on the 3’UTR region to create a 1.1 Kb homology region outside of guide RNA 3 and 2kb homology region outside of guide RNA 4. PAM sites were altered on the donor construct for guide RNA 3 (GGG->CCG)) and guide RNA 4 (CGG-> CAG). guide RNA 3 (GGGCTTACGCCGGTATTAG) and guide RNA 4 (GCTTTGGGTGTAGGAGCGTC) were cloned into a dual U6 (1+3 promoter) expression construct pCFD4 provided by the Mann lab using Gibson assembly (New England Biolabs). The donor construct and the guide RNA construct were injected into *wlig4; attp40 [nos-Cas9]/Cyo; ci-[w^+^] / ci-[w^+^]* embryos. Surviving adults were crossed to *yw hs-flp; Sp/Cyo; TM2/TM6B; Dp[y+]/Dp[y+]* flies. Male *crCi/Dp[y+]* “transformants” were identified by white eyes, amplified into suitable stocks and genotyped for sequences encoding Flag and HA tags upstream and downstream of *ci* coding sequence, respectively. Balanced *ci* alleles were further genotyped to confirm the mutation of interest.

### Donor Template Cloning

For crCi-WT, ΔCORD, P(1-3)A, S849A, S849AΔCORD, Δ760-934, Δ1270-1370, ΔCDN, ΔCDNΔCORD plasmid design was developed using APE software. Overlapping primer PCR reactions were used to add, mutate, and delete regions on Ci with PfuUltraII Fusion HS DNA polymerase (Agilent Technologies). PCR products were introduced into the Zero Blunt Topo cloning vector (Invitrogen). The alterations in Ci were then re-introduced from the Zero Blunt Topo cloning Vector into the BSK-F1 Donor construct using compatible enzymes or Gibson Assembly (New England Biolabs). The final constructs were fully sequenced (Genewiz).

### Drosophila stocks

Drosophila stocks were maintained on standard cornmeal/molasses/agar medium at room temperature.

Females of the genotype *yw hs-flp; ptc-lacZ/TM6B, Tb; ci^94^/Dp[y^+^]* were crossed to *yw hs-flp; Sp/Cyo; gCi-WT/ΔCORD/S849A; ci^94^/Dp[y^+^]* males, selecting third instar larval progeny lacking y^+^ and Tb to obtain wing discs with third chromosome transgenes as the only source of Ci.

Females of the *genotype yw hs-flp; ptc-lacZ/TM6B, Tb; ci^94^/Dp[y^+^]* were crossed to *yw hs-flp; Sp/Cyo; crCi-X/ Dp[y^+^]* males, selecting third instar larval progeny lacking *y^+^* and *Tb* to obtain wing discs with a single constructed *crCi* allele as the only source of Ci.

Females of the genotype *yw hs-flp; Su(fu)^LP^ ptc-lacZ/TM6B, Tb; ci^94^/Dp[y^+^]* were crossed to *yw hs-flp; Sp/Cyo; Su(fu)^LP^/TM6B, Tb; crCi-X/ Dp[y^+^]* males, selecting third instar larval progeny lacking *y^+^* and *Tb* to obtain wing discs with a single constructed *crCi* allele as the only source of Ci in a Su(fu) null background.

Females of the genotype (“2L”) *yw hs-flp UAS-GFP; tub-Gal80 FRT40A/Cyo; C765-GAL4 ptc-lacZ/TM6B, Tb; ci^94^/Dp[y^+^]* were crossed to males of the genotype *yw hs-flp; pka-C1^H2^ FRT40A/Cyo; gCi-WT/ΔCORD/S849A /TM6B, Tb; ci^94^/Dp[y^+^]* or *yw hs-flp; pka-C1^H2^ FRT40A/Cyo; crCi-X / Dp[y^+^]*, selecting third instar larval progeny lacking *y^+^* and *Tb* to obtain wing discs with a single constructed *crCi* allele as the only source of Ci and GFP-marked *pka* mutant clones.

Females of the genotype (“2b”) *yw hs-flp UAS-GFP; FRT42D P[Ci^+^] tub-Gal80 /Cyo; C765-GAL4 ptc-lacZ/TM6B, Tb; ci^94^/Dp[y^+^]* were crossed to males of the genotype *yw hs-flp; FRT42D/Cyo; crCi-X / Dp[y^+^]*, selecting third instar larval progeny lacking *y^+^* and *Tb* to obtain wing discs with a single constructed *crCi* allele as the only source of Ci in GFP-marked clones lacking *P[Ci^+^]* with neighboring cells including *P[Ci^+^]*.

Females of the genotype (“2b”) *yw hs-flp UAS-GFP; FRT42D P[Ci^+^] tub-Gal80 /Cyo; C765-GAL4 ptc-lacZ/TM6B, Tb; ci^94^/Dp[y^+^]* were crossed to males of the genotype *yw hs-flp; FRT42D cos2^2^/Cyo; gCi-WT/ΔCORD/S849A /TM6B, Tb; ci^94^/Dp[y^+^]* or *yw hs-flp; FRT42D cos2^2^/Cyo; crCi-X / Dp[y^+^]*, selecting third instar larval progeny lacking *y^+^* and *Tb* to obtain wing discs with a single constructed *crCi* allele as the only source of Ci in GFP-marked clones lacking *cos2* activity and *P[Ci^+^]* with neighboring cells expressing *P[Ci^+^]*.

Females of the genotype (“2b”) *yw hs-flp UAS-GFP; FRT42D P[Ci^+^] tub-Gal80 /Cyo; C765-GAL4 ptc-lacZ/TM6B, Tb; ci^94^/Dp[y^+^]* were crossed to males of the genotype *yw hs-flp; smo^2^FRT42D UAS-GAP-Fu/Cyo; crCi-X / Dp[y^+^]*, selecting third instar larval progeny lacking *y^+^* and *Tb* to obtain wing discs with a single constructed *crCi* allele as the only source of Ci in GFP-marked clones expressing GAP-Fu and lacking *P[Ci^+^]* with neighboring cells expressing *P[Ci^+^]*. Females of the genotype (“2R”) *yw hs-flp UAS-GFP; smo^2^ FRT42D P[Smo^+^] tub-Gal80 /Cyo; C765-GAL4 ptc-lacZ/TM6B, Tb; ci^94^/Dp[y^+^]* were crossed to males of the genotype *yw hs-flp; FRT42D cos2^2^/Cyo; crCi-X / Dp[y^+^]*, selecting third instar larval progeny lacking *y^+^* and *Tb* to obtain wing discs with a single constructed *crCi* allele as the only source of Ci and GFP-marked clones lacking *cos2* activity.

Females of the genotype (“2R”) *yw hs-flp UAS-GFP; smo^2^ FRT42D P[Smo^+^] tub-Gal80 /Cyo; C765-GAL4 ptc-lacZ/TM6B, Tb; ci^94^/Dp[y^+^]* were crossed to males of the genotype *yw hs-flp; smo^2^ FRT42D UAS-GAP-Fu/Cyo; crCi-X / Dp[y^+^]*, selecting third instar larval progeny lacking *y^+^* and *Tb* to obtain wing discs with a single constructed *crCi* allele as the only source of Ci in GFP-marked clones expressing GAP-Fu and lacking *smo* activity.

Females of the genotype *yw hs-flp fu^mH63^; FRT42D P[y^+^] P[Fu+]/Cyo; (Su(fu)^LP^) C765-GAL4 ptc-lacZ/TM6B, Tb; ci^94^/Dp[y^+^]* were crossed to males of the genotype *yw hs-flp; Sp/Cyo; (Su(fu)^LP^/TM6B); crCi-X / Dp[y^+^]*, selecting male third instar larval progeny lacking y^+^ and Tb to obtain wing discs lacking Fu kinase activity (with or without functional Su(fu)) and a single constructed *crCi* allele as the only source of Ci.

### Immunohistochemistry

Wing disc clones were generated by heat-shocking late first or early second instar larvae for 1h at 37°C and dissections took place 3.5 to 4 days later in wandering third instar larvae. Wing discs were dissected from late third instar larvae in PBS and fixed in 4% paraformaldehyde (in PBS) for 30 min, rinsed 3X with PBS, blocked with 10% normal goat serum (Jackson ImmunoResearch Laboratories, Inc.) in PBS-T (0.1% Triton) for 1h, and stained with the following primary antibodies: rabbit anti–β-galactosidase (1:10000; MP Biomedicals), mouse 4D9 anti-Engrailed (1:5 Developmental Studies Hybridoma Bank), Rat 2A1 anti-Ci (1:3 Developmental Studies Hybridoma Bank), overnight at 4°. Inverted Larvae were then washed three times in PBST for 10 min each and incubated with Alexa Fluor 488, 546, 594, or 647 secondary antibodies (1:1,000; Molecular Probes) for 1h at room temperature. Larvae were washed twice in PBST for 20 min each, once in PBS for 10 min and mounted in Aqua/Poly mount (Polysciences).

### Quantitation from fluorescent images

Fluorescence images were captured using 20x, 63x, or 40x (discs with far anterior clones) objectives using 1.4 NA oil immersion lenses on a confocal microscope (LSM 700 and LSM800; Carl Zeiss). The range indicator was used to set the appropriate laser intensity per experiment for each fluorophore such that the signal was in the linear range.

#### Intensity Profiles

To measure intensity profiles along the AP axis, an elongated rectangle was drawn on a central region of the wing pouch, avoiding the D/V border. The y-axis shows the average fluorescence intensity over the height of the rectangle at each point on the x-axis (AP axis) for *ptc-lacZ* expression or Ci-155 protein, measured using Image J software (NIH, Bethesda, Maryland). For 20x images, at least 3 wings discs per condition were measured and averaged for each plot, using the posterior edge of *ptc-lacZ* expression as a reference point for the AP border.

#### Clone Measurements

The average fluorescent intensity of *ptc-lacZ* or Ci-155 over specific regions was measured using Image J. Multiple clones or clone regions (for large clones), anterior regions (at least 3 per disc), AP border sections (at least 3 per disc), posterior regions (at least 3 per disc), were analyzed for each disc. To make sure the best region was acquired for measurements in clones the region was selected using the GFP marker in the central part of the clone and confirmed to not be on a fold or shadowed region. For the AP border, regions were measured avoiding the DV boundary and abnormal folds. For Figures 4 and 6, *ptc-lacZ* clone intensity was calculated relative to AP border levels after subtracting anterior cell intensity values from each because *ptc-lacZ* is sometimes expressed artifactually in posterior cells: (clone-averaged anterior)/(averaged AP border-averaged anterior). In Figure 5 *ptc-lacZ* intensity in clones and anterior cells outside clones was in each case divided by AP border intensity without any subtractions. Ci-155 clone intensity and intensity in anterior cells outside clones (Fig. 4) were calculated relative to AP border levels after subtracting posterior cell intensity values from each: (clone-averaged posterior)/(averaged AP border-averaged posterior) and (anterior-averaged posterior)/(averaged AP border-averaged posterior). For Figure 5 *ptc-lacZ* intensity measurements in clones and anterior regions were divided by the AP-Border.

### Adult Wings

Adult wings were pulled off anaesthetized flies and placed in 70% ethanol for 5 minutes, transferred to 100% ethanol, and then mounted in Aqua/Poly Mount (Polysciences). They were imaged with Transmitted Light on a Nikon Diaphot 300 microscope using a 10x objective.

