## Supplemental Figures for "Drosophila Hedgehog can act as a morphogen in the absence of regulated Ci processing"

**Supplementary Figure Legends**

**Figure S1. Loss of Hib binding sites does not greatly affect Ci-155 activity or Hh-stimulated proteolysis.**

(A-F) *ptc-lacZ* (red) and Ci-155 (black) intensity profiles for wing discs expressing one copy of the designated *ci* transgene or alleles from Fig. 2A-F. (G-K) *ptc-lacZ* (red) and (G'-K') Ci-155 (gray-scale) in wing discs with one copy of the indicated *crCi* alleles (with $ci^{94}$) and (J, K) homozygous loss of Su(fu) at high (63x objective) magnification, with AP boundary (dashed yellow line at posterior *ptc-lacZ* boundary). Scale bars are (G-K) 40μm. (L, M) Intensity profiles for (L) Ci-155 and (M) *ptc-lacZ* for *crCi-WT* and *crCi-S3-5* in the presence (black and red, respectively) or absence (gray and green, respectively) of Su(fu). Arrows in (L) indicate the locations of *ptc-lacZ* initial rise (orange), 50% increase (brown) and peak (green) for *crCi-WT* discs. (N, O) Anterior En (green) induction, revealed by *ptc-lacZ* (red) marking of the AP boundary (yellow line) extended further anterior for (N) *crCi-WT* than (O) *crCi-S3-5*. Scale bars 40μm. (P) *ptc-lacZ* intensity profiles (from Fig. 2G"-J") for *crCi-P(1-3)A* (green), *crCi-S849A* (red) and *crCi-WT* (black) were overlapping, while *ptc-lacZ* induction was much lower for *crCi-Δ1270-1370* (gray).

**Figure S2. Processing-resistant Ci-155 induces ectopic dpp expression and anterior disc expansions are suppressed by adding a constitutive Ci repressor.**

(A, B) Wing discs with *gCi-S849A* and no other source of functional Ci (B) induced ectopic anterior *dpp-lacZ* (gray-scale) in addition to the AP border stripe but (A) ectopic *dpp-lacZ* expression is suppressed by a wild-type *ci* allele (on the $Dp[y^{+}]$ chromosome). (C, D) Wing discs with one copy of (C) *gCi-WT* or (D) *gCi-S849A* have similar expression of *ptc-lacZ* (red) confined to the AP border (at a lower level than in wing discs without $ci^{Ce}$) and normal morphology in the presence of a *ci* allele that produces constitutive Ci repressor ($ci^{Ce}$). 2A1 antibody staining (gray-scale) is high throughout the anterior in both cases because the $ci^{Ce}$ product includes the 2A1 epitope and does not undergo regulated processing. (E, F) Wing discs with anterior clones (GFP, green, yellow arrows) that have lost a second chromosome *gCi* transgene, leaving one copy of the indicated *crCi* alleles as a source of Ci. *ptc-lacZ* was clearly detected in clones expressing only (E) CiΔ760-934 but not (F) in clones expressing CiΔ1270-1370 (arrows), albeit at lower levels than at the AP border (arrowheads). (G, H) Anterior En was (G) nearly absent for CiΔ760-934 and (H) much reduced for CiΔ1270-1370 in wing discs expressing one copy of each *crCi* allele as the only source of Ci. Scale bars (A-D) 100μm and (E, F) 40μm.

**Figure S3. Reduced AP border activity of Ci lacking the CORD domain.**

(A-H) Wing discs expressing one copy of *crCi-WT* or *crCi-ΔCORD* as the only source of Ci, showing (A, C, E, F) Ci-155 (gray-scale), (B, D, G, H, A'-H') *ptc-lacZ* (red) or (B, D, G, H, B", D", G", H") En (green) and AP compartment boundary (yellow line). Reduced En induction by Ci-ΔCORD was (B, D) not restored to normal by loss of Su(fu). (E-H) Loss of Fu kinase activity in $fu^{mH63}$ wing discs produced (E, F) similarly strong, broad Ci-155 at the AP border and (E', F') drastic reduction of *ptc-lacZ* for Ci-WT and Ci-ΔCORD, while (G, H) additional loss of Su(fu) restored *ptc-lacZ* without anterior En in both cases. (I-L) induction of *ptc-lacZ* (red) in *smo* mutant clones (green, arrows) expressing GAP-FU was similar for Ci-WT and Ci-ΔCORD encoded by either (I, J) *gCi* transgenes or (K, L) *crCi* alleles and was much lower than at the AP border (arrowheads) in all cases, as presented graphically in (N). Much higher *ptc-lacZ* (red), matching AP border levels (arrowheads) were observed in GAP-Fu clones (green, arrows) expressing processing-resistant Ci-S849AΔCORD, accompanied by (M") significant Ci-155 (gray-scale) proteolysis. Scale bars are (A, C, E, F) 100μm and (D, G, H, I-M) 40μm.

### Figure S1

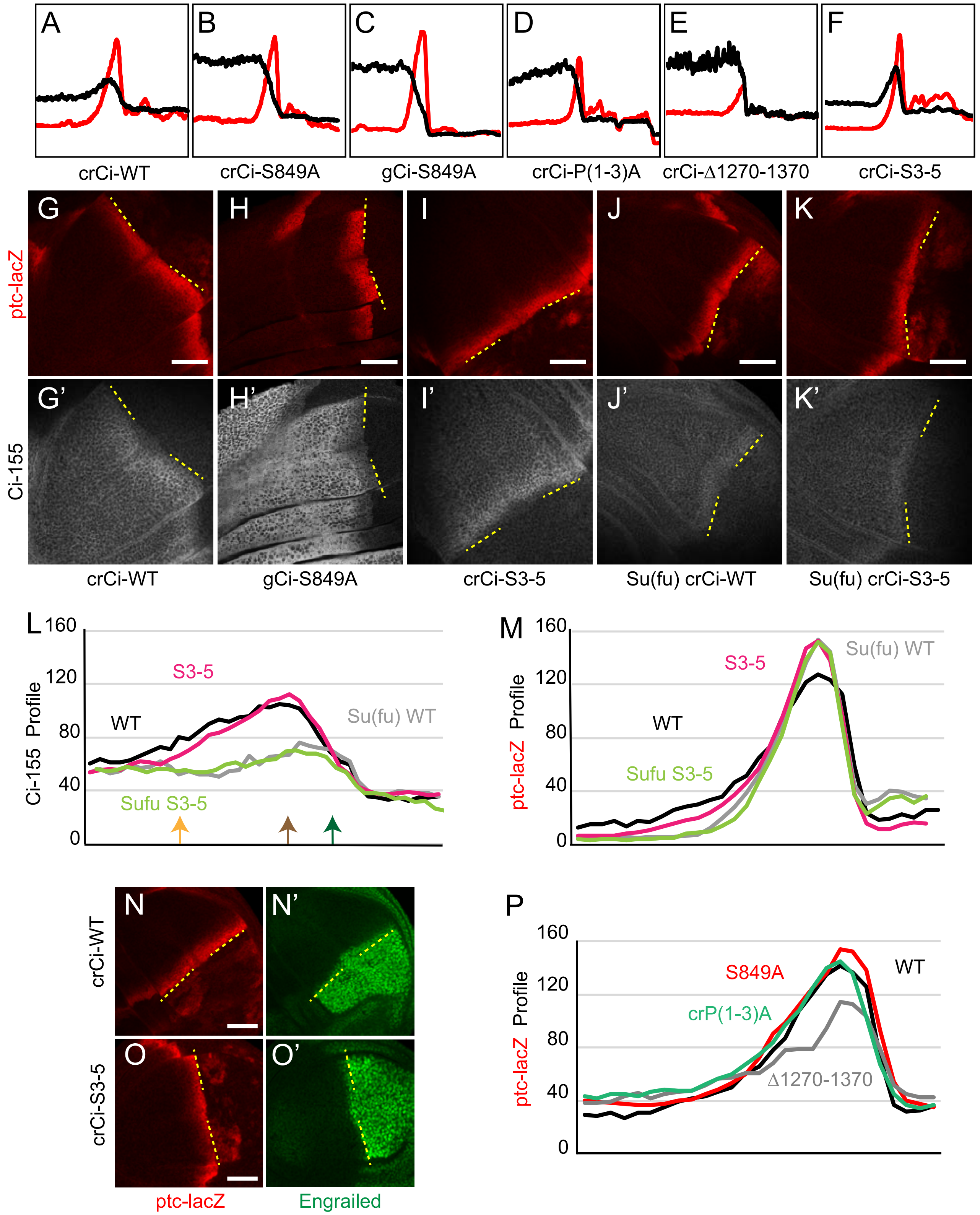

A
B
C
D
E
F
crCi-WT
crCi-S849A
gCi-S849A
crCi-P(1-3)A
crCi-Δ1270-1370
crCi-S3-5
G
H
I
J
K
ptc-lacZ
G'
H'
I'
J'
K'
Ci-155
crCi-WT
gCi-S849A
crCi-S3-5
Su(fu) crCi-WT
Su(fu) crCi-S3-5
L
Ci-155 Profile
160
120
80
40
0
S3-5
WT
Su(fu) WT
Sufu S3-5
M
ptc-lacZ Profile
160
120
80
40
0
Su(fu) WT
S3-5
WT
Sufu S3-5
N
N'
O
O'
crCi-WT
crCi-S3-5
ptc-lacZ
Engrailed
P
ptc-lacZ Profile
160
120
80
40
0
S849A
WT
crP(1-3)A
Δ1270-1370

### Figure S2

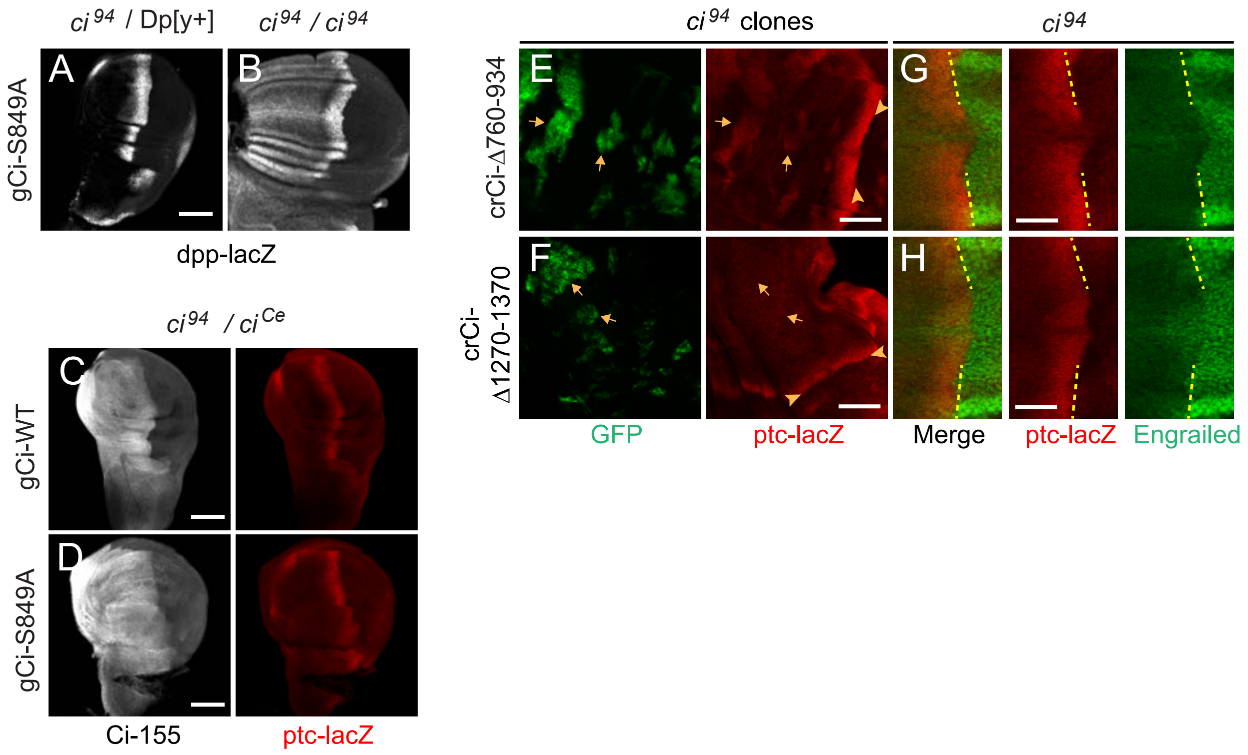

ci 94 / Dp[y+]
ci 94 / ci 94
gCi-S849A
A
B
dpp-lacZ
ci 94 / ci Ce
gCi-WT
C
gCi-S849A
D
Ci-155
ptc-lacZ
ci 94 clones
ci 94
crCi-Δ760-934
E
G
crCi-Δ1270-1370
F
H
GFP
ptc-lacZ
Merge
ptc-lacZ
Engrailed

### Figure S3

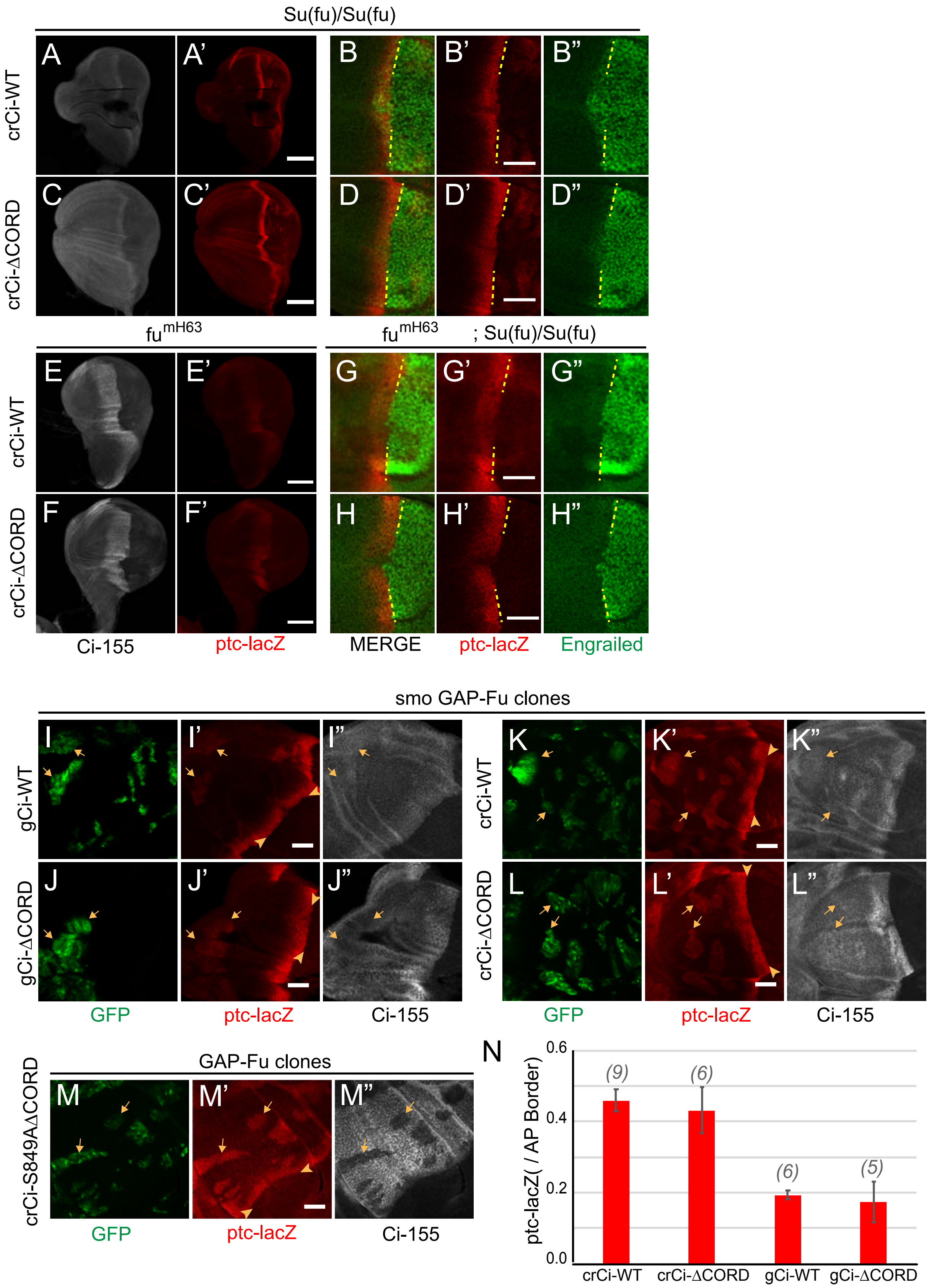
Su(fu)/Su(fu)
crCi-WT
crCi-ΔCORD
A
A'
B
B'
B"
C
C'
D
D'
D"
fu mH63
fu mH63 ; Su(fu)/Su(fu)
E
E'
F
F'
G
G'
G"
H
H'
H"
Ci-155
ptc-lacZ
MERGE
ptc-lacZ
Engrailed
smo GAP-Fu clones
gCi-WT
gCi-ΔCORD
I
I'
I"
J
J'
J"
crCi-WT
crCi-ΔCORD
K
K'
K"
L
L'
L"
GFP
ptc-lacZ
Ci-155
GFP
ptc-lacZ
Ci-155
crCi-S849AΔCORD
GAP-Fu clones
M
M'
M"
GFP
ptc-lacZ
Ci-155
N
ptc-lacZ( / AP Border)
0.6
0.4
0.2
0.0
(9)
(6)
(6)
(5)
crCi-WT
crCi-ΔCORD
gCi-WT
gCi-ΔCORD
